# Observing one-divalent-metal-ion dependent and histidine-promoted His-Me family I-PpoI nuclease catalysis in crystallo

**DOI:** 10.1101/2024.05.02.592236

**Authors:** Caleb Chang, Grace Zhou, Yang Gao

## Abstract

Metal-ion-dependent nucleases play crucial roles in cellular defense and biotechnological applications. Time-resolved crystallography has resolved catalytic details of metal-ion-dependent DNA hydrolysis and synthesis, uncovering the essential roles of multiple metal ions during catalysis. The histidine-metal (His-Me) superfamily nucleases are renowned for binding one divalent metal ion and requiring a conserved histidine to promote catalysis. Many His-Me family nucleases, including homing endonucleases and Cas9 nuclease, have been adapted for biotechnological and biomedical applications. However, it remains unclear how the single metal ion in His-Me nucleases, together with the histidine, promotes water deprotonation, nucleophilic attack, and phosphodiester bond breakage. By observing DNA hydrolysis *in crystallo* with His-Me I-PpoI nuclease as a model system, we proved that only one divalent metal ion is required during its catalysis. Moreover, we uncovered several possible deprotonation pathways for the nucleophilic water. Interestingly, binding of the single metal ion and water deprotonation are concerted during catalysis. Our results reveal catalytic details of His-Me nucleases, which is distinct from multi-metal-ion-dependent DNA polymerases and nucleases.

## Introduction

Mg^2+^-dependent nucleases play fundamental roles in DNA replication and repair ^1–4^, RNA processing ^5–8^, as well as immune response and defense ^9–11^. Moreover, they are widely employed for genome editing in biotechnological and biomedical applications ^12,13^. These nucleases are proposed to cleave DNA through a SN_2_ type reaction, in which a water molecule, or sometimes, a tyrosine side chain ^14^, initiates the nucleophilic attack on the scissile phosphate with the help of metal ions ^15^. Metal ions can orient and stabilize the binding of the negatively charged nucleic acid backbone ^16^, promoting proton transfer, nucleophilic attack, and stabilization of the transition state. As a highly varied family of enzymes, Mg^2+^-dependent nucleases can be broadly categorized by the number of metal ions captured in their active site. So far, Mg^2+^-dependent nucleases with one and two metal ions have been observed. In the two-Mg^2+^-ion dependent nuclease, a metal ion binds on the leaving group side of the scissile phosphate (Me^2+^_B_) while the other binds on the nucleophile side (Me^2+^_A_). In one-metal ion dependent nucleases, only the metal ion corresponding to the B site in two-metal ion dependent nucleases is present ^17^. Moreover, transiently bound metal ions have been identified in the previously thought two-metal-ion RNaseH ^18^ and EndoV nucleases ^19^ *via* time-resolved X-ray crystallography, proposed to play key roles in various stages of their catalysis. Similarly, catalysis in the one-metal ion dependent APE1 nuclease has been observed *in crystallo*, but mechanistic details regarding its metal-ion dependence have not been thoroughly explored ^20–22^. There also exist three-Zn^2+^ dependent nucleases, with two Zn^2+^ binding in the A-equivalent and B-equivalent positions, while the third Zn^2+^ coordinating the sp oxygen on the nucleophile side of the scissile phosphate ^23^. However, it remains unclear how the single metal ion in one-metal-ion dependent nucleases is capable of aligning the substrate, promoting deprotonation and nucleophilic attack, and stabilizing the pentacovalent transition state.

A large subfamily of one-metal-ion dependent nucleases consist of histidine-metal (His-Me) nucleases that perform critical tasks in biological pathways such as apoptosis ^24,25^, extracellular defense ^26,27^, intracellular immunity (CRISPR-Cas9) ^28,29^, and intron homing (homing endonuclease) ^30,31^. Despite sharing poor sequence homology, the structural cores and active sites of His-Me nucleases are highly conserved and thus are proposed to catalyze DNA hydrolysis through a similar mechanism ^15,31,32^. Surrounded by two beta sheets and an alpha helix, the His-Me nucleases active sites all contain a single metal ion, a histidine, and an asparagine (Fig. 1A, S1A, and S2). The asparagine residue helps to stabilize metal ion binding whereas the strictly conserved histidine is suggested to deprotonate a nearby water for nucleophilic attack towards the scissile phosphate ^33–36^. To this day, the dynamic reaction process of DNA hydrolysis by His-Me nucleases has never been visualized, and the mechanism of single metal ion-dependent and histidine-promoted catalysis remains unclear. Emerging genome editing and disease treatment involving CRISPR-Cas9 emphasize the importance of understanding the catalytic mechanism of DNA hydrolysis by His-Me nucleases ^37–39^. Recent crystal and cryo-electron microscope structures of Cas9 have captured the His-Me family Cas9 histidine-asparagine-histidine (HNH) active site engaged with DNA before and after cleavage ^40–43^. However, due to the relatively low resolution and the static nature of the structures, key catalytic details, such as metal ion dependence, transition-state stabilization, and alternative deprotonation pathways, remain elusive. Moreover, the large size and multiple conformational checkpoints during Cas9 catalysis ^40–43^ hinder *in crystallo* observation of Cas9 catalysis.

**Fig. 1.**
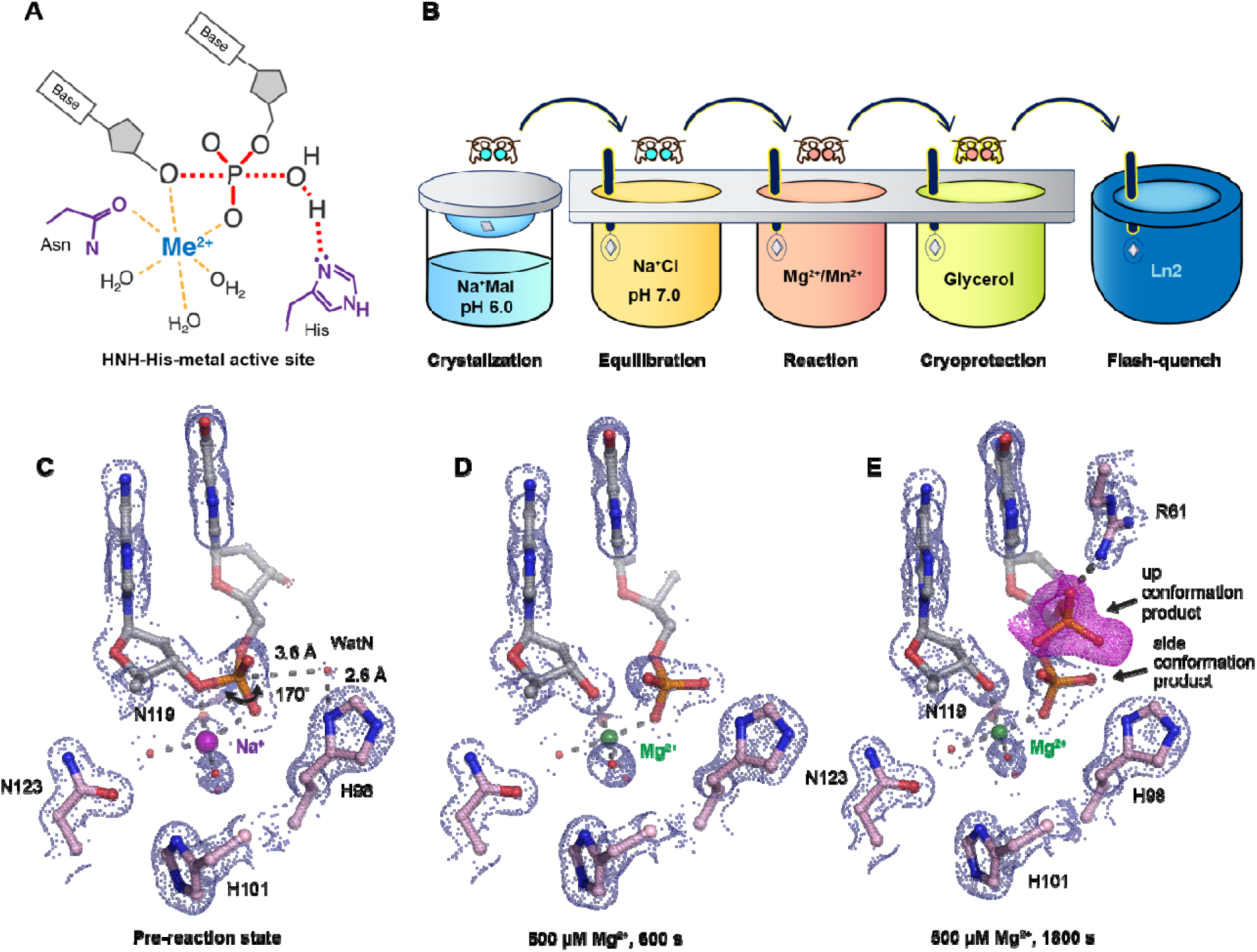
Observing His-Me family I-PpoI catalyze DNA hydrolysis *in crystallo*. (**A**) Model of one-metal-ion dependent and histidine promoted His-Me enzyme catalysis and transition state stabilization. (**B**) Metal ion soaking setup for *in crystallo* observation of DNA hydrolysis with I-PpoI. (**C-E**) Structural intermediates of I-PpoI *in crystallo* DNA cleavage showcasing the pre-reaction state in (**C**) and product states in (**D**) and (**E**). The 2F_o_-F_c_ map for Me^2+^, DNA, waters (red spheres), and catalytic residues (blue) was contoured at 2.0 σ (σ values represent r.m.s. density values). (**E**) The F_o_-F_c_ omit map for the up conformation of the product (violet) was contoured at 3.0 σ.

I-PpoI is a well-characterized intron-encoded homing endonuclease member of the *physarum polycephalum* slimemold. By 1999, Stoddard and colleagues were able to capture intermediate structures of I-PpoI complexed with DNA before and after product formation ^33,44^. We herein employed I-PpoI as a model system and applied time-resolved crystallography to observe the catalytic process of His-Me nuclease. By determining over 40 atomic resolution structures of I-PpoI during its reaction process, we show that one and only one divalent metal ion is involved in DNA hydrolysis. Moreover, we uncover several possible deprotonation pathways for the nucleophilic water. Notably, metal ion binding and water deprotonation are highly concerted during catalysis. Our findings provide mechanistic insights into one-metal-ion dependent nucleases, enhancing future design and engineering of these enzymes for emerging biomedical applications.

## Results

### Preparation of the I-PpoI system for in crystallo DNA hydrolysis

We sought to implement I-PpoI for *in crystallo* metal ion soaking (Fig. 1B), which has been successful in elucidating the catalytic mechanisms of DNA polymerases ^45–50^, nucleases ^18–21^, and glycosylase ^51^. First, a complex of I-PpoI and a palindromic DNA was crystalized at pH 6 with 0.2 M sodium malonate. Similar to previous studies, a dimer of I-PpoI was found in the asymmetric unit, with both active sites engaged for catalysis and DNA in the middle bent by 55° (Fig. S1A). The two I-PpoI molecules were almost identical and thus served as internal controls for evaluating the reaction process (Fig. S1B). Within each I-PpoI active site, a water molecule (nucleophilic water, WatN) existed 3.6 Å from the scissile phosphate, near the imidazole side chain of His98. On the leaving group side of the scissile phosphate, the metal exhibited an octahedral geometry, being coordinated by three water molecules, two oxygen atoms from the scissile phosphate, and the conserved asparagine (Fig. 1C).

Next, we removed malonate in the crystallization buffer, which may chelate metal ions and hinder metal ion diffusion ^52^, by equilibrating the crystals in 200 mM NaCl buffer at pH 6, 7, or 8 for 30 min. The diffraction quality of the crystals was not affected during the soaking. The structures of I-PpoI equilibrated at pH 6, 7, or 8 in the presence of 200 mM NaCl showed no signs of product formation and appeared similar to the malonate and previous reported I-PpoI structures ^33,44^ (Fig. S3A). To confirm if a monovalent metal ion binds in the pre-reaction state (PRS), we soaked the I-PpoI crystals in buffer containing Tl^+^ and detected anomalous electron density at the metal ion binding site without product formation (Fig. 3SB). Our results support that the monovalent metal ion can bind within the active site without initiating reaction, in corroboration with our biochemical assays (Fig. S3C).

### Witnessing DNA hydrolysis in crystallo by I-PpoI

To initiate the chemical reaction *in crystallo*, we transferred the I-PpoI crystals equilibrated in NaCl buffer to a reaction buffer with 500 µM Mg^2+^ (Fig. 1B). After 600 s soaking, we saw a significant negative F_o_-F_c_ peak on the leaving group side of the scissile phosphate atom as well as a positive F_o_-F_c_ peak on the other side (Fig. S3D), indicating that DNA hydrolysis was occurring *in crystallo*. After DNA cleavage, the newly generated phosphate group shifted 1 Å towards the WatN (Fig. 1D). Furthermore, an additional product state was observed after soaking the crystals in 500 µM Mg^2+^ for 600 s and 1800 s, at which the newly formed phosphate shifted 3 Å away from the metal ion to form a hydrogen bond with Arg61 (Fig. 1E). Our results confirm that the implementation of I-PpoI with *in crystallo* Me^2+^ soaking is feasible for observing the I-PpoI catalytic process and dissecting the mechanism of His-Me nucleases.

With an established *in crystallo* reaction system, we next monitored the reaction process by soaking I-PpoI crystals in buffer containing 500 µM Mg^2+^ pH 7, for 10-1200 s. Density between the reactant phosphate and nucleophilic water increased along with longer soaking time, indicating the generation of phosphate products (Fig. 2A). During the reaction, the coordination distances of the metal ion ligands decreased 0.1-0.2 Å when Mg^2+^ exchanged with Na^+^, which is consistent with their preferred geometry ^53^ (Fig. 2B, C). At reaction time 160 s, 45% of product had been generated (Fig. 2D), which later plateaued to 65% at 300 s. During the reaction process, the sugar ring of the reactant and product DNA that resides around 3 Å away from the scissile phosphate, remained in a C3’-endo conformation. As Mg^2+^ soaking time increased from 10-160 s, we observed the WatN approaching the scissile phosphate (Fig. 2E). At the same time, the conserved His98 sidechain proposed to deprotonate the WatN was slightly rotated.

**Fig. 2.**
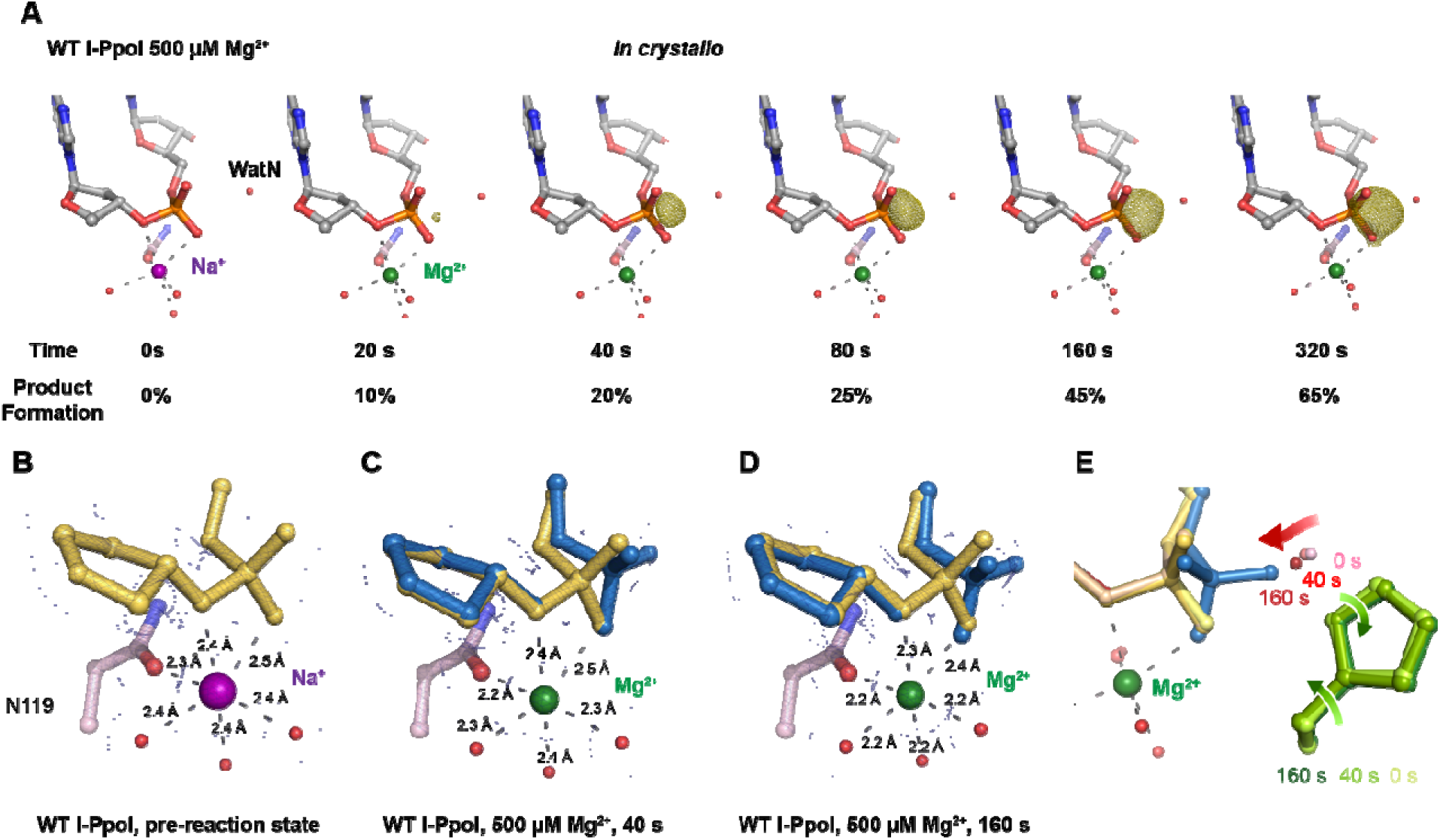
*In crystallo* DNA hydrolysis by I-PpoI. (**A**) Structures of I-PpoI during *in crystallo* catalysis after 500 µM Mg^2+^ soaking for 0 s, 20 s, 40 s, 80 s, 160 s, 320 s. The F_o_-F_c_ omit map for the product phosphate (green) was contoured at 3.0 σ. (**B-D**) I-PpoI complexes featuring metal ion coordination when bound with Na^+^ in (**B**) Mg^2+^ in the earlier time point of the reaction process in (**C**), and Mg^2+^ in the later time point of the reaction process in (**D**). The 2F_o_-F_c_ map for Me^2+^, DNA, waters (red spheres), and catalytic residues (blue) was contoured at 2.0 σ. (**E**) Alignment of the WatN and rotation in His98 during I- PpoI reaction.

**Fig. 3.**
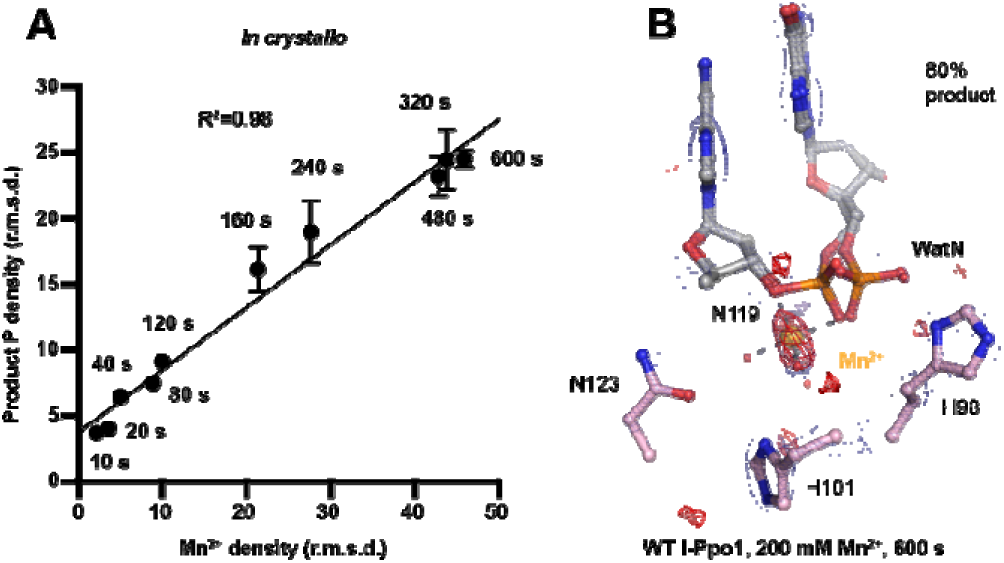
Detection of Me^2+^ binding during DNA hydrolysis *in crystallo*. (**A**) Correlation (R^2^) between the newly formed phosphate and Mn^2+^ binding at pH 6. The points represent the mean of duplicate measurements for the electron density of the reaction product phosphate within two I-PpoI molecules in the asymmetric unit while the errors bars represent the standard deviation. (**B**) Additional Mn^2+^ binding sites were not detected after 0.9765 Å X-ray diffraction in the I-PpoI active site after 10 min soaking in 200 mM Mn^2+^. The 2F_o_-F_c_ map for Me^2+^, DNA, waters (red spheres), and catalytic residues (blue) was contoured at 2.0 σ. The anomalous map for Mn^2+^ was contoured at 3.0 σ.

### A single divalent metal ion was captured during DNA hydrolysis

Throughout the reaction process with Mg^2+^, we found that the electron density for the Mg^2+^ metal ion strongly correlated (R^2^=0.97) with product formation (Fig. S4A, C), which may suggest that saturation of this metal ion site is required and sufficient for catalysis. However, Mg^2+^’s similar size to Na^+^ makes it suboptimal for quantifying metal ion binding. To thoroughly investigate metal-ion dependence, we repeated the *in crystallo* soaking experiment with Mn^2+^, which is more electron rich and can be unambiguously assigned based on its electron density and anomalous signal. With 500 µM Mn^2+^ in the reaction buffer, we found that the reaction process and the product conformation were similar to that for Mg^2+^. After 160 s Mn^2+^ soaking, clear anomalous signal was present at the metal ion binding site, confirming the binding of Mn^2+^ (Fig. S4F). For crystal structures of I-PpoI with partial product formation, Mn^2+^ signal at the metal ion site correlated with product formation with a R^2^ of 0.98 (Fig. 3A and S4B). To further search if additional and transiently-bound divalent ions participate in the reaction, we soaked the I-PpoI crystals in high concentration of Mn^2+^ (200 mM) for 600 s, at which 80% product formed within the active site. However, apart from the single metal ion binding site, we do not detect anomalous signal for additional Mn^2+^, despite such high concentration of Mn^2+^ (Fig. 3B and S5). The strong correlation between product phosphate formation with Mn^2+^ binding and the absence of additional anomalous density peaks in heavy Mn^2+^ soaked crystals suggest that one and only one divalent metal ion is involved in I-PpoI DNA hydrolysis.

To examine if additional monovalent metal ions participate during DNA hydrolysis, we titrated Tl^+^, which can replace monovalent Na^+^ or K^+^ and yields anomalous signal ^54,55^, at up to 100 mM concentration in the biochemical assay. Apart from precipitation that occurred at 50 and 100 mM Tl^+^, we found that product conversion remained unaffected with increasing Tl^+^ in solution (Fig. S4G). Similarly, increasing Na^+^ concentration in solution did not increase product conversion (Fig. S4H). In addition, only one Tl^+^ was detected by its anomalous signal in the pre-reaction state (Fig. S3B). The biochemical and structural results indicate that additional monovalent metal ion may not be necessary for DNA hydrolysis. However, crystal deterioration limited us from soaking the I-PpoI crystals in high concentration of Tl^+^ for *in crystallo* reaction.

### pH-dependence of I-PpoI DNA cleavage

During DNA hydrolysis, deprotonation of the WatN is required for the nucleophilic attack and phosphodiester bond breakage. This proton transfer has been proposed to be mediated by the highly conserved His98 ^33,44^ that lies within 3 Å from the WatN. We speculated that the ability of His98 to activate the nearby WatN and mediate proton transfer would be affected by pH. The DNA cleavage assay revealed that I-PpoI cleavage activity increased with pH with a pKa of 8.3 (Fig. 4A), which is much higher than the pKa of histidine, but the histidine pKa and the proton transfer process may be affected by the active site environment. To explore how pH affects the active site configuration and catalysis, we conducted *in crystallo* soaking experiments with Mg^2+^ at pH 6 and pH 8 in addition to the pH 7 data series in Figure 2. Consistent with in solution experiments, higher pH resulted in faster product formation *in crystallo* (Fig. 4B), indicating that metal ion binding may also be affected by pH. To test this, we performed Mg^2+^ titration at different pH in solution. Our results showed that over 100-times higher concentration of Mg^2+^ was needed to yield 50% product in pH 6 versus pH 8 (Fig. 4C), confirming that pH affects metal-ion dependent I-PpoI catalysis. Likewise, the reactions *in* crystallo at higher pH reached 50% product formation at shorter soaking times (320 s at pH 6, 160 s at pH 7, and 80 s at pH 8, respectively). Interestingly, the crystal structures that contained 30-35% product at pH 6 and pH 8 were nearly identical (Fig. 4D, E). The positions of the His98 residue and the WatN were practically superimposable (Fig. 4F). At all pH, Mg^2+^ binding strongly correlated with product formation (R^2^ greater than 0.95), suggesting that low pH reduces the overall reaction rate without altering the reaction pathway (Fig. S4D, E).

**Fig. 4.**
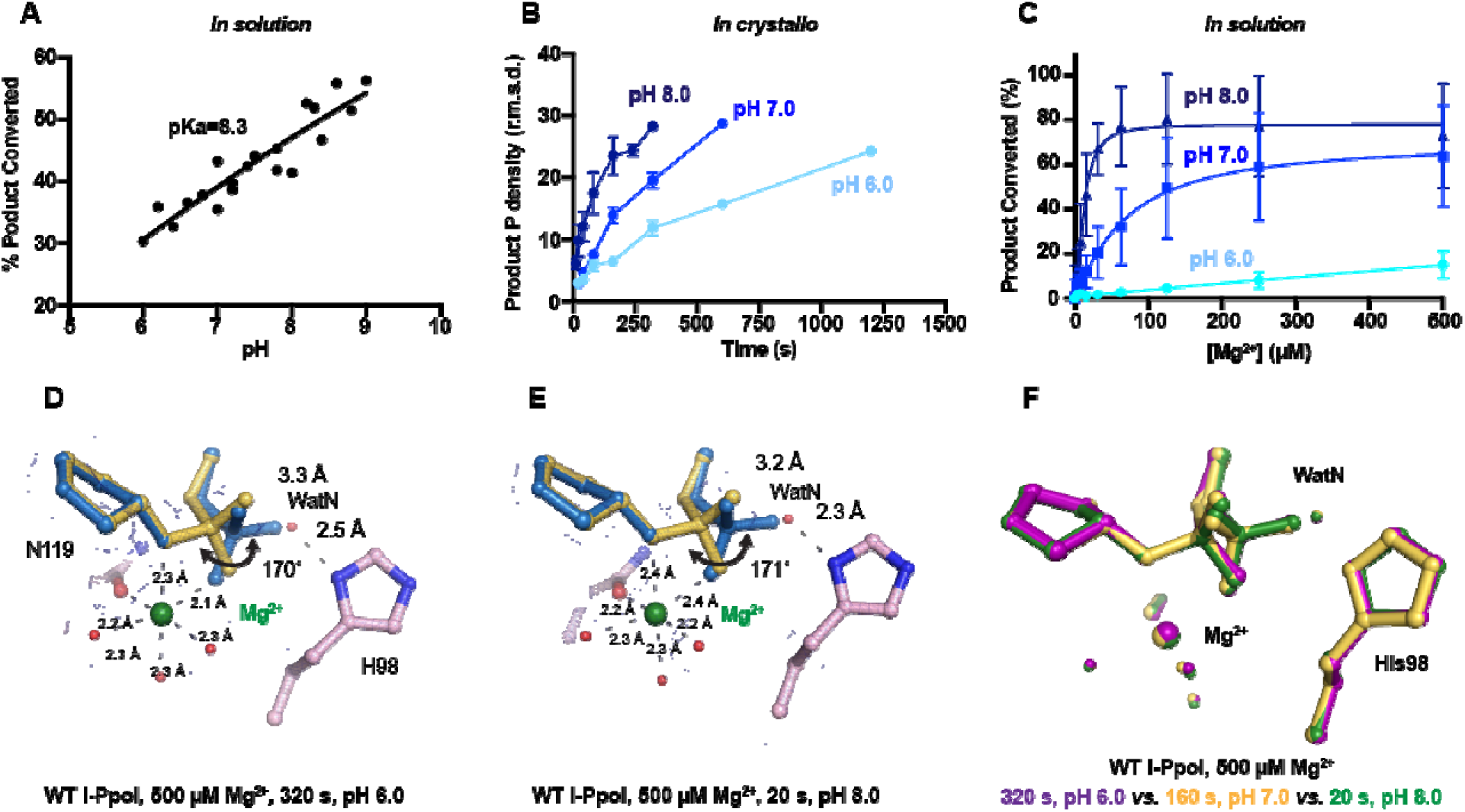
Effect of pH on I-PpoI DNA hydrolysis. (**A**) DNA hydrolysis by WT I-PpoI with increasing pH in solution. (**B**) DNA hydrolysis by WT I-PpoI *in crystallo* at pH 6, 7, and 8. The points represent the mean of duplicate measurements for the electron density of the reaction product phosphate after a period of Mg^2+^ soaking within two I-PpoI molecules in the asymmetric unit. The errors bars represent the standard deviation. (**C**) The effect of pH on metal ion dependence in solution. The points represent the mean of triplicate measurements for the percentage of cleaved DNA product while the errors bars represent the standard deviation. (**D**) Structure of I-PpoI *in crystallo* DNA hydrolysis at pH 6 after 320 s of 500 µM Mg^2+^ soaking. (**E**) Structure of I-PpoI *in crystallo* DNA hydrolysis at pH 8 after 20 s of 500 µM Mg^2+^ soaking. (**D, E**) The 2F_o_-F_c_ map for Me^2+^, DNA, waters (red spheres), and catalytic residues (blue) was contoured at 2.0 σ. (**F**) Structural comparison of the active site after 500 µM Mg^2+^ soaking for 320 s at pH 6 (purple), 160 s at pH 7 (yellow), and 20 s at pH 8 (green).

### Nucleophilic water deprotonation pathway during I-PpoI DNA cleavage

We next investigated the deprotonation pathway with mutagenesis. Because His98 has been proposed to primarily activate the nucleophilic water, we first mutated His98 to alanine (Fig. 5A). As expected, cleavage activity of H98A I-PpoI drastically dropped. Our assays showed that H98A I-PpoI displayed residual activity but required a reaction time of 1 hr to be comparable to WT I-PpoI (Fig. 5B), similar to the H98Q mutant in previous studies ^56^. Furthermore, varying the pH resulted in a sigmoidal activity curve of I-PpoI pH dependence, corresponding to a pKa of 7.9 (Fig. 5C). The significantly reduced reaction rate and the shift of pH dependence suggest that His98 plays a key role in pH sensing and water deprotonation. On the other hand, the low but existing activity of H98A I-PpoI suggests the presence of alternative general bases for proton transfer. Another histidine (His78) resides on the nucleophile side 3.8 Å from the WatN, with a cluster of water molecules in between (Fig. 5A). We hypothesized that His78 may substitute His98 as the proton acceptor. The DNA cleavage assays revealed that the single mutant, H78A I-PpoI, had an activity similar to WT, whereas the double mutant (H78A/H98A I-PpoI) exhibited much lower activity than that of H98A (Fig. 5B). Furthermore, varying the pH for H78A/H98A I-PpoI resulted in a sigmoidal activity curve that corresponded to a pKa of 8.7 (Fig. 5C). In consistent with previous speculations, our results confirm the possibility of His78 as an alternative general base ^56^. The residual activity and pH dependence of H78A/H98A I-PpoI indicated that something else was still activating the nucleophilic water in the absence of any nearby histidine. Furthermore, we found that titrating imidazole in H98A and H78A/H98A I-PpoI partially rescued cleavage activity (Fig. S6A, B), similar as observed in Cas9 and EndA nuclease ^57,58^. Collectively, our results indicate that His98 is the primary proton acceptor like previous simulation results ^59^ but at the same time, I-PpoI can use alternative pathways to activate the nucleophilic WatN.

**Fig. 5.**
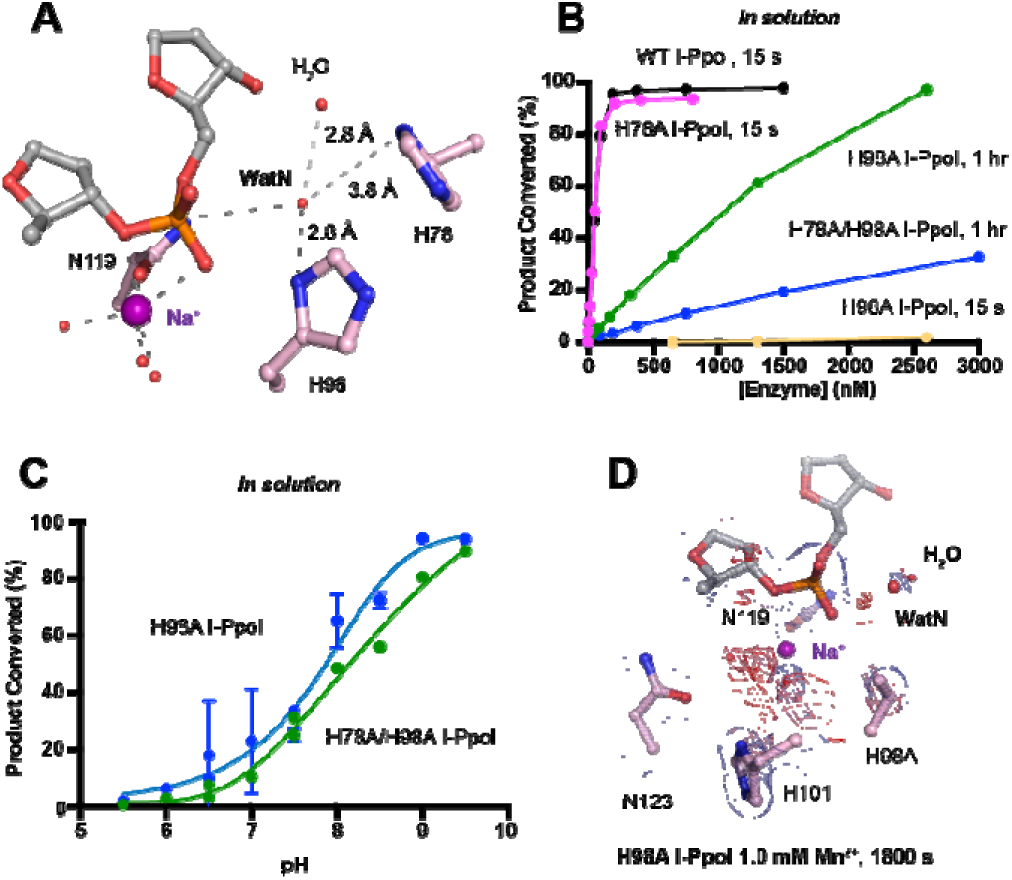
Nucleophilic water deprotonation during I-PpoI DNA hydrolysis. (**A**) Active site environment surrounding WatN. His78 exists near the WatN besides His98. (**B**) In solution DNA hydrolysis activity of various I-PpoI histidine mutants. The points represent the mean of triplicate measurements for the percentage of cleaved reaction product while the errors bars (too small to see) represent the standard deviation. (**C**) DNA hydrolysis by H98A I-PpoI (blue) and H78A/H98A I-PpoI (green) at various pH in solution. (**D**) Structure of H98A I-PpoI active site after 0.9765 Å X-ray diffraction after 1 mM Mn^2+^ soaking for 1800 s. The anomalous map for Mn^2+^ was contoured at 2.0 σ. The 2F_o_-F_c_ map for Me^2+^, DNA, waters (red spheres), and catalytic residues (blue) was contoured at 2.0 σ. (**B, C**) The points represent the mean of triplicate measurements for the percentage of generated reaction product while the errors bars represent the standard deviation.

We next sought to understand the mechanism of His98-promoted hydrolysis from a structural standpoint. In our *in crystallo* soaking experiments, we equilibrated H98A I-PpoI crystals in 500 µM Mg^2+^. The structure looked nearly identical to the Na^+^ structure and previous H98A I-PpoI structure with Mg^2+^. As shown earlier, the coordination environment of Na^+^ and Mg^2+^ was quite similar. To confirm divalent metal ion binding, the H98A I-PpoI crystals were soaked for 1800 s in 500 µM Mn^2+^, the same concentration used to initiate DNA hydrolysis by WT I-PpoI. However, to our surprise, the metal ion binding site was devoid of any anomalous signal. Increasing the Mn^2+^ concentration to 1 mM still did not produce anomalous density at the Me^2+^ binding site (Fig. 5D and S6C). The results indicate that metal ion binding can be altered by perturbing His98 and possibly water deprotonation, even though the metal ion binding site and His98 exist 7 Å apart without direct interaction. Although we tried soaking the H98A I-PpoI crystals in 1 mM Mg/Mn^2+^ and 100 mM imidazole for 15 h, metal ion binding, imidazole binding or product formation were not detected (Fig. S6D), possibly due to the difficulty of imidazole diffusion within the lattice. Our results indicate that perturbing the deprotonation pathway not only affects nucleophilic attack but also metal binding, suggesting that I-PpoI catalyzes DNA catalysis *via* a concerted mechanism.

## Discussion

Time-resolved crystallography can visualize time-dependent structural changes and elucidate mechanisms of enzyme catalysis with unparalleled detail *in crystallo*, especially for light-dependent enzymes, in which the reactions can be synchronously initiated by light pulses ^60–63^. In complementary, recently advanced metal ion diffusion-based time-resolved crystallographic techniques have uncovered rich dynamics at the active site and transient metal ion binding during the catalytic processes of metal-ion-dependent DNA polymerases ^45–50^, nucleases ^18,19^ and glycosylase ^51^. In addition to metal ions captured in static structures, these transient metal ions have been shown to play critical roles in catalysis, such as water deprotonation for nucleophilic attack in RNaseH ^18^, bond breakage and product stabilization in polymerases ^45–48,50,64,65^ and alignment of the substrate and nucleophilic water in MutT ^66^. Interestingly, one and only one metal ion was captured within the I-PpoI active site during catalysis, even when high concentration (200 mM Mn^2+^) of metal ion was tested (Fig. 3). The observed location of the Me^2+^ on the leaving group side of the scissile phosphate corresponds to the Me^2+^_B_ in two metal-ion dependent nucleases ^17^ (Fig. S2). However, the metal ion is unique in its environment and role. First, it is coordinated by a water cluster and asparagine side chain (sometimes with an additional aspartate residue) rather than the acidic aspartate and glutamic acid clusters that outline the active sites of RNaseH and APE1 nuclease ^67^ as well as DNA polymerases (Fig. S7). Even in the absence of divalent Mg^2+^ or Mn^2+^, the active site including the scissile phosphate was already well-aligned in I-PpoI, which is again different from RNaseH. Instead, Leu116 and the beta sheet consisting of Arg61, Gln63, Lys65, Thr67 ^44,68^ helped to position the DNA optimally towards the active site (Fig. S1A, C). Second, single metal ion binding is strictly correlated with product formation in all conditions, at different pH and with different mutants (Fig. 3A and S4A, B, C-E)^58^. Thus, similar to the third metal ion in DNA polymerases and RNaseH, the metal ion in I-PpoI is not required for substrate alignment but is essential for catalysis. We suspect that the single metal ion helps stabilize the transition state and reduce the electronegative buildup of DNA, thereby promoting DNA hydrolysis.

Proton transfer by a general base is essential for a SN_2_-type nucleophilic attack. Such deprotonation of the nucleophilic water has been attributed to His98, which is highly conserved in His-Me nucleases. Existing close to the nucleophilic water at 2.6 Å, His98 is perfectly positioned to mediate the proton transfer. Moreover, due to the bulky presence of His98 and beta sheet protein residues, there is no space for an additional metal ion at the nucleophilic side (Fig. S5). The H98A mutation significantly reduced catalytic activity and altered pH-dependence. But since H98A I-PpoI showed residual activity, His78, imidazole, or its surrounding waters may still serve as alternative general bases for accepting the proton, similar to Pol η, in which primer 3’-OH deprotonation can occur through multiple pathways ^50^. However, the order of events regarding metal ion binding, water deprotonation, and nucleophilic attack remains unanswered. Based on our *in crystallo* observations, water deprotonation and metal ion binding appeared to be highly correlated (Fig 6A). Lowering the pH not only reduced reaction rate but also slowed metal ion binding (Fig. 4B, C). Moreover, the metal ion was not observed *in crystallo* when His98 was removed (Fig. 5D). As there is no direct interaction between His98 and the Me^2+^ binding site, the divalent metal ion may be sensitive to the charge potential of the substrate scissile phosphate, which may be indirectly affected by the deprotonated state of the nucleophilic water. Conversely, binding of the divalent metal ion may alter the local electrostatic environment and affect His98 deprotonation. Consistently, previous molecular dynamics simulation of Cas9 has suggested that the histidine pKa is highly sensitive to active site changes ^69^. Without a proper proton acceptor, the metal ion may be prone for dissociation without the reaction proceeding, and thus stable Mg^2+^ binding was not observed *in crystallo* without His98 (Fig. 6B). On the other hand, optimal alignment of the metal ion and WatN within the active site, labeled as metal-binding state, leads to irreversible bond breakage (Fig. 6A). In summary, our experimental observations suggest a concerted mechanism for one-metal-ion promoted DNA hydrolysis, offering guidance for future computational analysis of enzyme catalysis ^69^ and the rational design and engineering of nucleases ^70^ for biotechnological and biomedical applications.

**Fig. 6.**
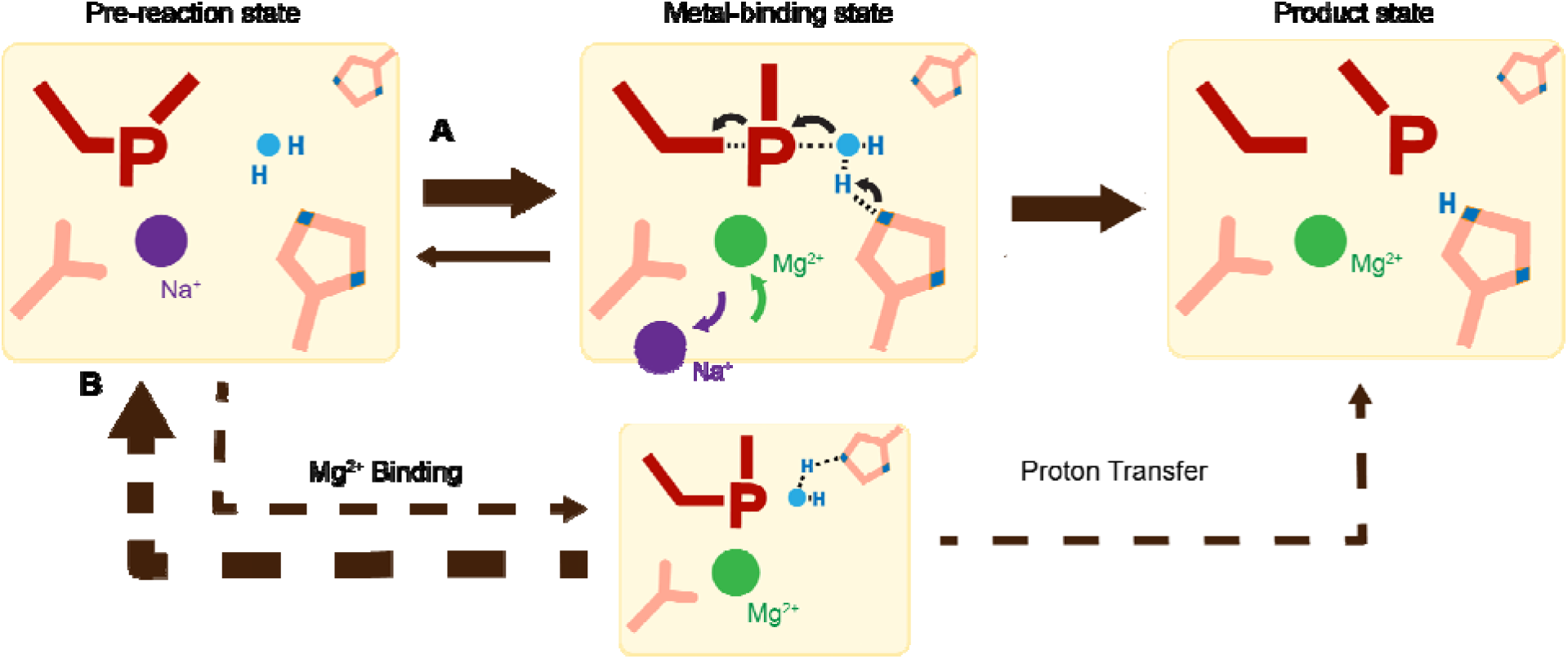
Catalytic model of His-Me nuclease DNA hydrolysis. Proposed model of His-Me nuclease DNA hydrolysis in which Me^2+^ binding, proton transfer, and nucleophilic attack are concerted (solid arrow) in the presence of the primary proton acceptor in (**A**) versus unfavored (dashed arrows) in the absence of the primary proton acceptor in (**B**).

## Materials and Methods

### Protein expression and purification

WT, His78Ala, His98Ala, H78A/H98A *physarum polycephalum* I-PpoI (residues 1-162) were cloned into a modified pET28p vector with a N-terminal 6-histidine tag and a PreScission Protease cleavage site. For protein expression, this I-PpoI plasmid was transformed into BL21 DE3 *E. coli* cells, which were grown in a buffer that contained (10 g/L glucose, 40 g/L α-lactose, 10% glycerol) for 24 hours (20°C). The cell paste was collected *via* centrifugation and re- suspended in a buffer that contained 50 mM Tris (pH 7.5), 1 M NaCl, 1mM MgCl_2_, 10 mM imidazole, 2 mM ß-mercaptoethanol (BME), and 5% glycerol. After sonification, I-PpoI was loaded onto a HisTrap HP column (GE Healthcare), which was pre-equilibrated with a buffer that contained 50 mM Tris (pH 7.5), 1 M NaCl, 1mM MgCl_2_, 10 mM imidazole, 2 mM ß- mercaptoethanol (BME), and 5% glycerol. The column was washed with 300 mL of buffer to remove non-specific bound proteins and was eluted with buffer that contained 50 mM Tris (pH 7.5), 1 M NaCl, 1mM MgCl_2_, 300 mM imidazole, and 2 mM ß-mercaptoethanol (BME). The eluted I-PpoI was incubated with PreScission Protease to cleave the N-terminal 6-histidine-tag. Afterwards, I-PpoI was desalted to 50 mM Tris (pH 7.5), 167 mM NaCl, 1mM MgCl_2_, 2 mM ß-mercaptoethanol (BME), and 5% glycerol and was loaded onto a Heparin column (GE Healthcare) equilibrated with 50 mM Tris (pH 7.5) and 167 mM NaCl. The protein was eluted with an increasing salt (NaCl) gradient, concentrated, and stored at 40% glycerol at -80°C.

### DNA hydrolysis assays

DNA hydrolysis activity of varying time was assayed by the following: The reaction mixture contained 100 nM WT I-PpoI, 50 mM NaCl, 100 mM Tris (pH7.5), 1.5 mM DTT, 0.05 mg/mL BSA, 50 nM DNA, 10 µM EDTA, and 4% glycerol. The hydrolysis assays were executed using a palindromic 5’-fluorescein- labelled DNA duplex (5’-TTG ACT CTC TTA AGA GAG TCA-3’). Reactions were initiated by adding 10 mM MgCl_2_ to the reaction mixture for 0-1 hr at 37 °C and were stopped by mixing with equal volume of a quench buffer, which contain 80% formamide, 100 mM EDTA (pH 8.0), 0.2 mg/ml xylene cyanol, and 0.2 mg/ml bromophenol.

The DNA hydrolysis activity at different pH was assayed by the following: The reaction mixture contained 100-3000 nM WT, H98A, and H78A/H98A I-PpoI, 50 mM NaCl, 1.5 mM DTT, 0.05 mg/mL BSA, 50 nM DNA, 10 µM EDTA, and 4% glycerol. Reactions were initiated by adding 50 mM MES (pH5.5-6.5), 50 mM HEPES (pH6.5-7.5), or 50 mM Tris (pH7.5-9.5) together with 10 mM MgCl_2_ to the reaction mixture for 30 min at 37 °C and were stopped by adding equal volume of quench buffer.

The DNA hydrolysis activity with different mutants was assayed by the following: The reaction mixture contained 50 mM NaCl, 100 mM Tris (pH7.5), 1.5 mM DTT, 0.05 mg/mL BSA, 50 nM DNA, 10 mM MgCl_2_, 10 µM EDTA, and 4% glycerol. Reactions were initiated by adding 0-3000 nM of WT, H78A, H98A, H78A/H98A I-PpoI to the reaction mixture for 15 s or 1 h at 37 °C and stopped by adding equal volume of quench buffer.

The DNA hydrolysis activity with different metal ions was assayed by the following: The reaction mixture contained 100 nM WT I-PpoI, 50 mM NaCl, 100 mM Tris (pH7.5), 1.5 mM DTT, 0.05 mg/mL BSA, 50 nM DNA, 10 µM EDTA, and 4% glycerol. Reactions were initiated by adding 10 mM of MgCl_2_, MnCl_2_, CaCl_2_, NiCl_2_, and ZnCl_2_ to the reaction mixture for 15 s at 37 °C and were stopped by adding equal volume of quench buffer.

The DNA hydrolysis activity with Tl^+^ or additional Na^+^ was assayed by the following: The reaction mixture contained 100-3000 nM WT I-PpoI, 100 mM Tris (pH7.5), 1.5 mM DTT, 0.05 mg/mL BSA, 50 nM DNA, 10 µM EDTA, and 4% glycerol. Reactions were conducted at 37 °C for 30 min by adding 0-350 mM TlCl or NaCl together with 10 mM MgCl_2_ to the reaction mixture and were stopped by adding equal volume of quench buffer.

The DNA hydrolysis activity with imidazole was assayed by the following: The reaction mixture contained 100-3000 nM H98A and H78A/H98A I-PpoI, 50 mM NaCl, 100 mM Tris (pH7.5), 1.5 mM DTT, 0.05 mg/mL BSA, 50 nM DNA, 10 µM EDTA, and 4% glycerol. Reactions were conducted at 37 °C for 30 min by adding 0-100 mM imidazole together with 10 mM MgCl_2_ to the reaction mixture and were stopped by adding equal volume of quench buffer.

For all reactions, after heating the quenched reaction mix to 97 °C for 5 min and immediately placing on ice, reaction products were resolved on 22.5% polyacrylamide urea gels. The gels were visualized by a Sapphire Biomolecular Imager and quantified using the built-in software. Quantification of percentage cleaved and graphic representation were executed by Graph Prism.

### Crystallization

WT or H98A I-PpoI in a buffer containing 20 mM Tris 7.5, 300 mM NaCl, 3 mM DTT, and 0.1 mM EDTA was added with (5’-TTG ACT CTC TTA AGA GAG TCA-3’) DNA at a molar molar of 1:1.5 for I-PpoI and DNA and added with 3-folds volume of buffer that contained 20 mM Tris 7.5, 3 mM DTT, and 0.1 mM EDTA. This I-PpoI-DNA complex was then cleaned with a Superdex 200 10/300 GL column (GE Healthcare) with a buffer that contained 20 mM Tris 7.5, 150 mM NaCl, 3 mM DTT, and 0.1 mM EDTA. The I-PpoI-DNA complex was concentrated to 2.8 mg/mL I-PpoI (confirmed by Bradford assay). All crystals were obtained using the hanging-drop vapour-diffusion method against a reservoir solution containing 0.1 M MES (pH 6.0), 0.2 M sodium malonate, and 20% (w/v) PEG3350 at room temperature within 4 days.

To identify the monovalent Me^+^ species that binds during the pre-reaction state, WT I-PpoI crystals were transferred and incubated in a buffer containing 0.1M MES (pH 6.0), 70 mM thallium acetate and 20% (w/v) PEG3350 for 30 min. Afterwards, the crystals were quickly dipped in a cryo-solution supplemented with 20% (w/v) glycerol and flash- cooled in liquid nitrogen.

### Chemical reaction *in crystallo*

The WT I-PpoI crystals were first transferred and incubated in a pre-reaction buffer containing 0.1M MES (pH 6.0 or 7.0) or 0.1M Tris (pH 8.0), 0.2 M NaCl, and 20% (w/v) PEG3350 for 30 min. The chemical reaction was initiated by transferring the crystals into a reaction buffer containing 0.1M MES (pH 6.0 or 7.0) or 0.1M Tris (pH 8.0), 0.2 M NaCl, and 20% (w/v) PEG3350, and 500 µM MgCl_2_ or MnCl_2_. After incubation for a desired time period, the crystals were quickly dipped in a cryo-solution supplemented with 20% (w/v) glycerol and flash- cooled in liquid nitrogen.

To observe any additional Me^2+^ binding sites during DNA hydrolysis, WT I-PpoI crystals were first transferred and incubated in a pre-reaction buffer containing 0.1M MES (pH 6.0), 0.2 M NaCl, and 20% (w/v) PEG3350 for 30 min. The chemical reaction was initiated by transferring the crystals into a reaction buffer containing 0.1M MES (pH 6.0), 0.2 M NaCl, and 20% (w/v) PEG3350, and 200 mM MnCl_2_. After incubation for 600 s, the crystals were quickly dipped in a cryo-solution supplemented with 20% (w/v) glycerol and flash- cooled in liquid nitrogen.

The metal ion soaking experiments with His98Ala I-PpoI were performed following the similar protocol as that of WT I-PpoI. His98Ala I-PpoI crystals were first incubated in a pre-reaction buffer containing 0.1M MES 7.0, 0.2 M NaCl, and 20% (w/v) PEG3350 for 30 min, followed by 1800s incubation in a reaction buffer containing 0.1M MES 7.0, 0.2 M NaCl, and 20% (w/v) PEG3350, and 1 mM MnCl_2_. To observe whether soaking in imidazole can initiate the reaction in the absence of His98, His98Ala I-PpoI crystals were first transferred and incubated in a buffer containing 0.1M MES (pH 6.0), 0.2 M NaCl, 1 mM MnCl_2_, and 20% (w/v) PEG3350 for 30 min. The crystals were then transferred into a reaction buffer containing 0.1M MES (pH 6.0), 0.2 M NaCl, 1 mM MgCl_2_ or MnCl_2_, and 100 mM imidazole, and 20% (w/v) PEG3350. After incubation for a desired time period, the crystals were quickly dipped in a cryo-solution supplemented with 20% (w/v) glycerol and flash- cooled in liquid nitrogen.

### Data collection and Refinement

Diffraction data were collected at 100 K on LS-CAT beam lines 21-D-D, 21-ID-F, and 21-ID-G at 1.1 Å or 0.97 Å at the Advanced Photon Source (Argonne National Laboratory) or beamlines 5.0.3. at 0.97 Å at ALS. Data were indexed in space group P3_1_21, scaled with XSCALE and reduced using XDS ^71^. Isomorphous I-PpoI structures with Na^+^ PDB ID 1CZ0 was used as initial models for refinement using PHENIX ^72^ and COOT ^73^.

Occupancies were assigned for the reaction product until there were no significant F_o_-F_c_ peaks. Occupancies were assigned for the metal ions, following the previous protocol ^46^ until 1) there were no significant F_o_-F_c_ peaks, 2) the B value had roughly similar values to its ligand 3) it matched the occupancy of the reaction product. For the structures in which some F_o_-F_c_ peaks were present around the Me^2+^ binding sites or reaction product, no change in the assigned occupancy was executed when a 10% change in occupancy (e.g. 100% to 90%) failed to significantly change the intensity of the F_o_-F_c_ peaks. Source data of the electron densities in r.m.s. density are provided as a Source Data file. Each structure was refined to the highest resolution data collected, which ranged between 1.42-2.2 Å. Software applications used in this project were compiled and configured by SBGrid ^74^. Source data of data collection and refinement statistics are summarized in **Table S1A-E**. All structural figures were drawn using PyMOL (http://www.pymol.org).

### Calculation of electron density

Electron density (r.m.s.d) from the F_o_-F_c_ map of the product phosphate and Me^2+^ were calculated by running a round of B-factor refinement in PHENIX after omitting the reaction product phosphate and Me^2+^ atoms from phase calculation in COOT. All structures from the same experiment (Mg^2+^-pH 6.0, Mg^2+^-pH 7.0, Mg^2+^-pH 8.0, Mn^2+^-pH 6.0) were refined to the lowest resolution in the same experiment group (1.80 Å for Mg^2+^-pH 6.0, 1.59 Å for Mg^2+^-pH 7.0, 1.70 Å for Mg^2+^-pH 8.0, and 1.80 Å for Mn^2+^-pH 6.0).

## Acknowledgments

Our sincere appreciation to the members of the Gao lab, and Drs. Phillips, Nikonowicz, and Lu who serve on CC’s thesis committee. We thank the APS LS-CAT beam technicians and research scientists Drs. Anderson, Wawrzak, Brunzelle, and Focia. This research used resources of the Advanced Photon Source, a U.S. Department of Energy (DOE) Office of Science User Facility operated for the DOE Office of Science by Argonne National Laboratory under Contract No. DE-AC02-06CH11357. Use of the LS-CAT Sector 21 was supported by the Michigan Economic Development Corporation and the Michigan Technology Tri-Corridor (Grant 085P1000817). The Berkeley Center for Structural Biology is supported in part by the Howard Hughes Medical Institute. The Advanced Light Source is a Department of Energy Office of Science User Facility under Contract No. DE-AC02-05CH11231. The Pilatus detector on 5.0.1. was funded under NIH grant S10OD021832. The ALS-ENABLE beamlines are supported in part by the National Institutes of Health, National Institute of General Medical Sciences, grant P30 GM124169.

## Author contributions

YG conceived the project. CC carried out all time-resolved crystallography experiments, data collection and processing. CC carried out protein purification, and CC and GZ executed protein crystallization. CC and GZ performed the biochemical assays. CC and YG wrote the manuscript.

## Data and materials availability

The coordinates, density maps, and structure factors for all the structures have been deposited in Protein Data Bank (PDB) under accession codes: 8VMO, 8VMP, 8VMQ, 8VMR, 8VMS, 8VMT, 8VMU, 8VMV, 8VMW, 8VMX, 8VMY, 8VMZ, 8VN0, 8VN1, 8VN2, 8VN3, 8VN4, 8VN5, 8VN6, 8VN7, 8VN8, 8VN9, 8VNA, 8VNB, 8VNC, 8VND, 8VNE, 8VNF, 8VNG, 8VNH, 8VNJ, 8VNK, 8VNL, 8VNM, 8VNN, 8VNO, 8VNP, 8VNQ, 8VNR, 8VNS, 8VNT, 8VNU. All data are available in the main text or the supplementary materials.

---

**Fig. S1.**
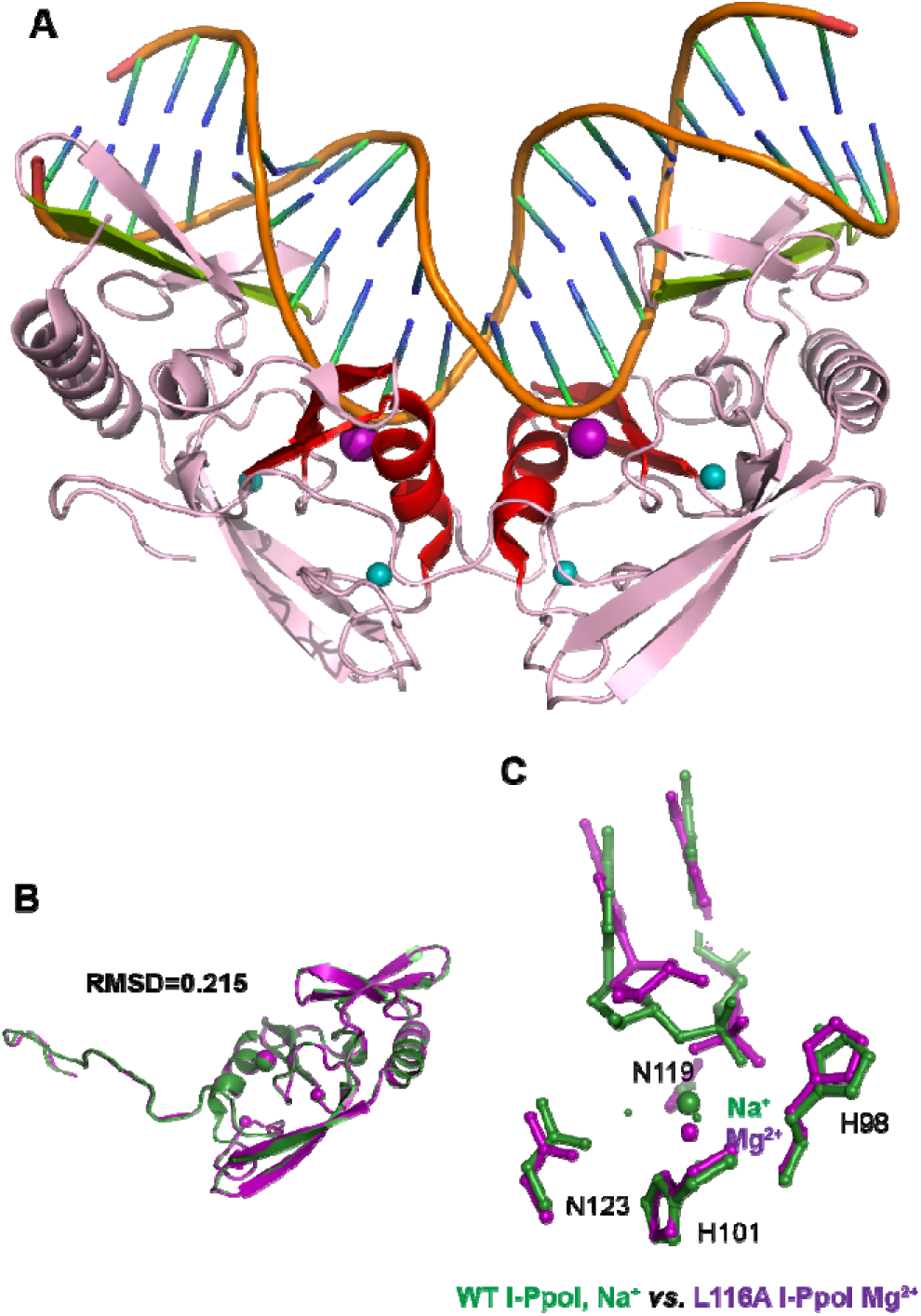
Overall structure and catalytic core of homing endonuclease I-PpoI. (**A**) Homing endonuclease I-PpoI (PDB ID 1CZO) binding as a dimer to bend DNA at 55°. The overall endonuclease is colored in pink while the DNA is colored in orange. The catalytic core comprised of an alpha helix and two beta sheets are highlighted in red. The metal ion binding site is depicted as a purple sphere while the Zn^2+^ binding sites are depicted as turquoise spheres. The beta sheets involved in DNA binding are depicted in light green. (**B**) Monomer of homing endonuclease I-PpoI (PDB ID 1CZO) (green) superimposed on top of the other I-PpoI monomer (purple) within the same unit cell, resulting in a RMSD of 0.215. (**C)**, Structural comparison of the active site of WT I-PpoI (PDB ID 1CZ0) versus Leu116Ala I-PpoI (PDB ID 1EVW). The DNA fails to dock tightly towards the Mg^2+^ in the active site of Leu116Ala I-PpoI.

**Fig. S2.**
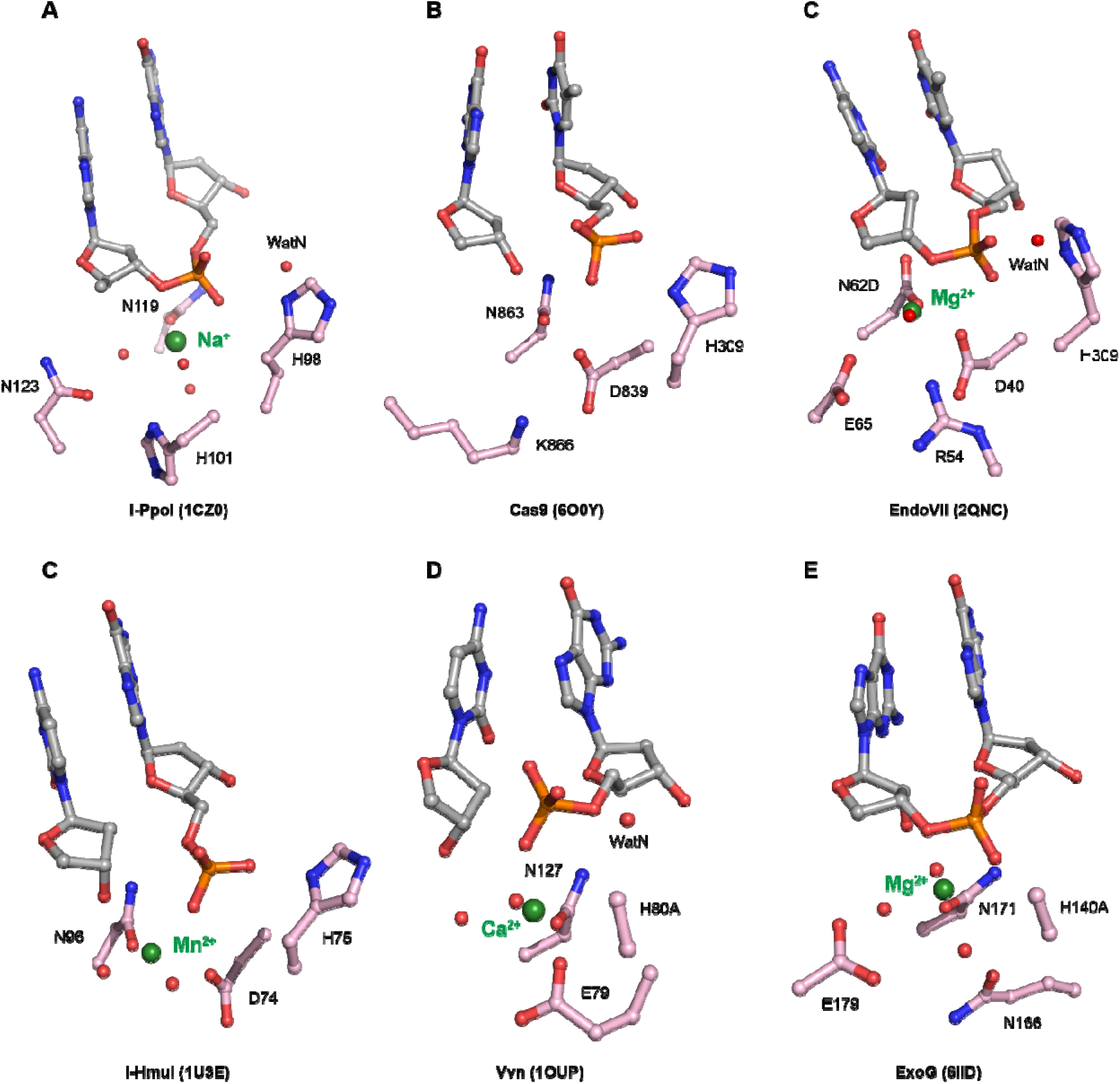
Active sites of His-Me superfamily nucleases. Carbon atoms of residues within the active site are colored in pink. The metal ions are depicted by green spheres while waters are depicted by red spheres.

**Fig. S3.**
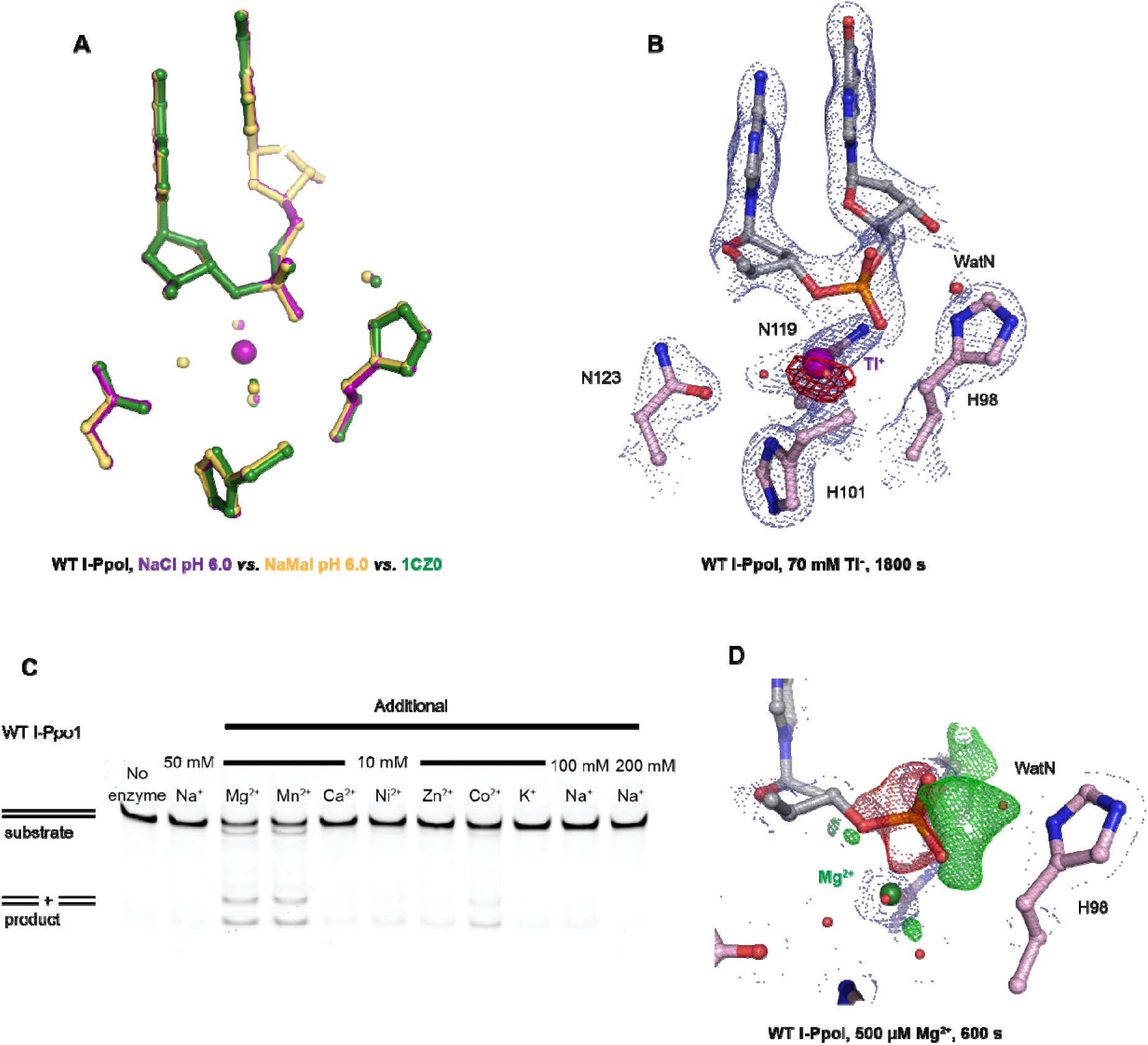
Establishing I-PpoI for *in crystallo* studies. (**A**) Structural comparison of the active site after soaking in NaCl in purple, sodium malonate in yellow and PDB ID 1CZ0 in green. (**B**) Tl^+^ anomalous signal was detected after 0.9765 Å X-ray diffraction at the I-PpoI metal ion binding site after 1800 s soaking in 70 mM Tl^+^. The 2F_o_-F_c_ map for Me^2+^, DNA, waters (red spheres), and catalytic residues (blue) was contoured at 2.0 σ. The anomalous map for Tl^+^ was contoured at 3.0 σ. (**C**) *In crystallo* metal ion assay of 10 mM additional metal ions on I-PpoI DNA hydrolysis. (**D**) Negative F_o_-F_c_ peaks (red) were detected on the leaving group side of the scissile phosphate while positive F_o_-F_c_ peaks (green) were detected on the nucleophile side after 600 s Mg^2+^ soaking. The 2F_o_-F_c_ map for Me^2+^, DNA, waters (red spheres), and catalytic residues (blue) was contoured at 2.0 σ. The negative F_o_-F_c_ map for the reactant phosphate (red) and the positive F_o_-F_c_ map for the product phosphate (green) were contoured at 3.0 σ.

**Fig. S4.**
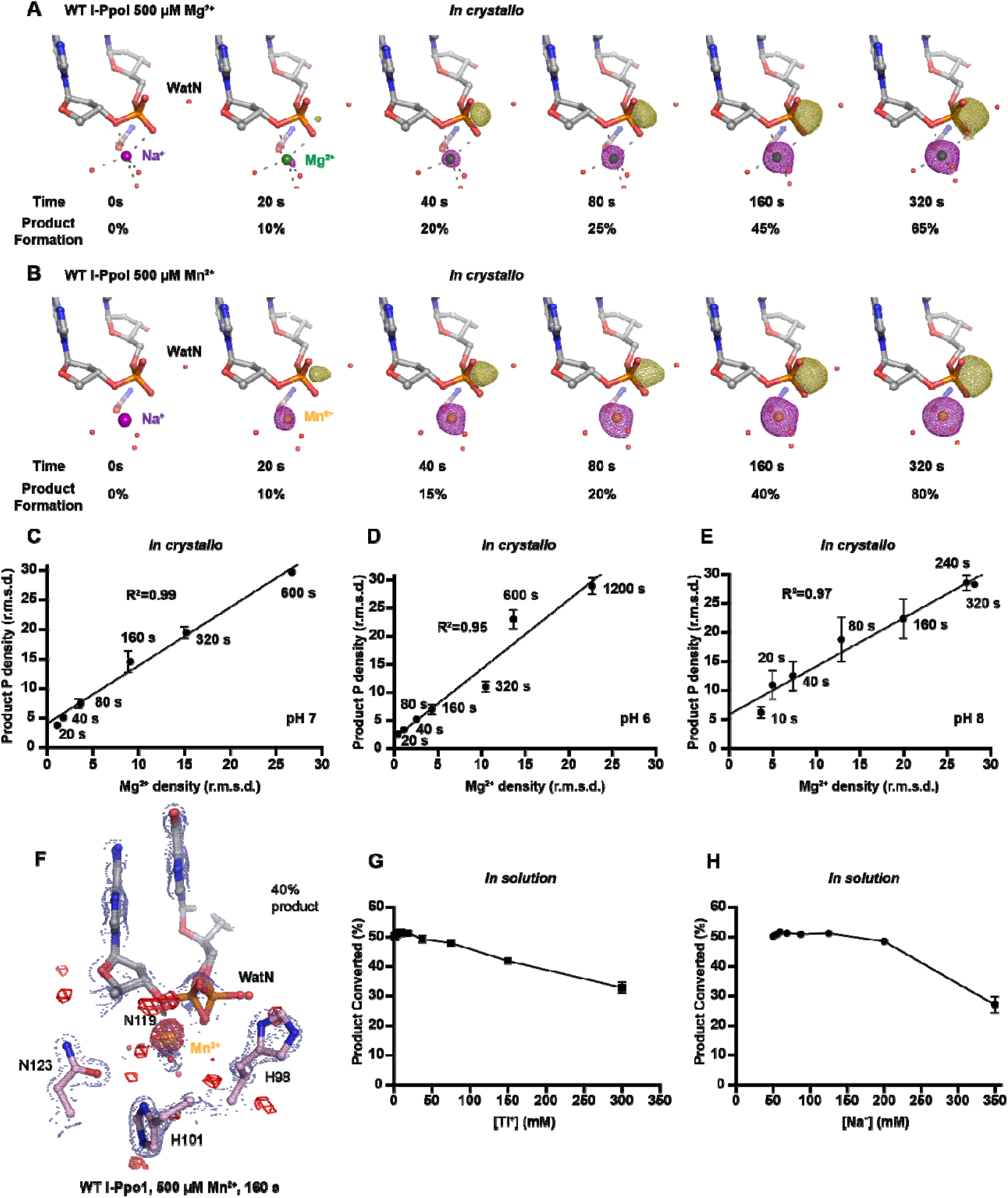
Additional metal ions are not required for DNA hydrolysis by I-PpoI. **(A**) Structures of I-PpoI during *in crystallo* catalysis after 500 µM Mg^2+^ soaking for 0 s, 20 s, 40 s, 80 s, 160 s, 320 s. The F_o_-F_c_ omit maps for the product phosphate (green mesh) and Mg^2+^ (purple mesh) were contoured at 3.0 σ. (**B**) Structures of I-PpoI during *in crystallo* catalysis after 500 µM Mn^2+^ soaking for 0 s, 20 s, 40 s, 80 s, 160 s, 320 s. The F_o_-F_c_ omit maps for the product phosphate (green mesh) and Mn^2+^ (purple mesh) were contoured at 3.0 σ. Correlation (R^2^) between the newly formed phosphate and Mg^2+^ binding *in crystallo* at pH 7 in (**C)**, pH 6 in (**D)**, and pH 8 in (**E**). (**C-E)** The points represent the mean of duplicate measurements for the electron density of the reaction product phosphate within two I-PpoI molecules in the asymmetric unit while the errors bars represent the standard deviation. (**F**) Mn^2+^ binding during DNA hydrolysis as revealed by anomalous signal of Mn^2+^ after 0.9786 Å X-ray diffraction. The 2F_o_-F_c_ map for Me^2+^, DNA, waters (red spheres), and catalytic residues (blue) was contoured at 2.0 σ. The anomalous map for Mn^2+^ was contoured at 3.0 σ. **(G)** Tl^+^ concentration on in solution DNA cleavage. Precipitation was detected when Tl^+^ was at or greater than 150 mM. **(H)** Additional Na^+^ concentration on in solution DNA cleavage. (**G, H**) The errors bars represent the standard deviation while the points represent the mean of triplicate measurements for cleaved DNA product.

**Fig. S5.**
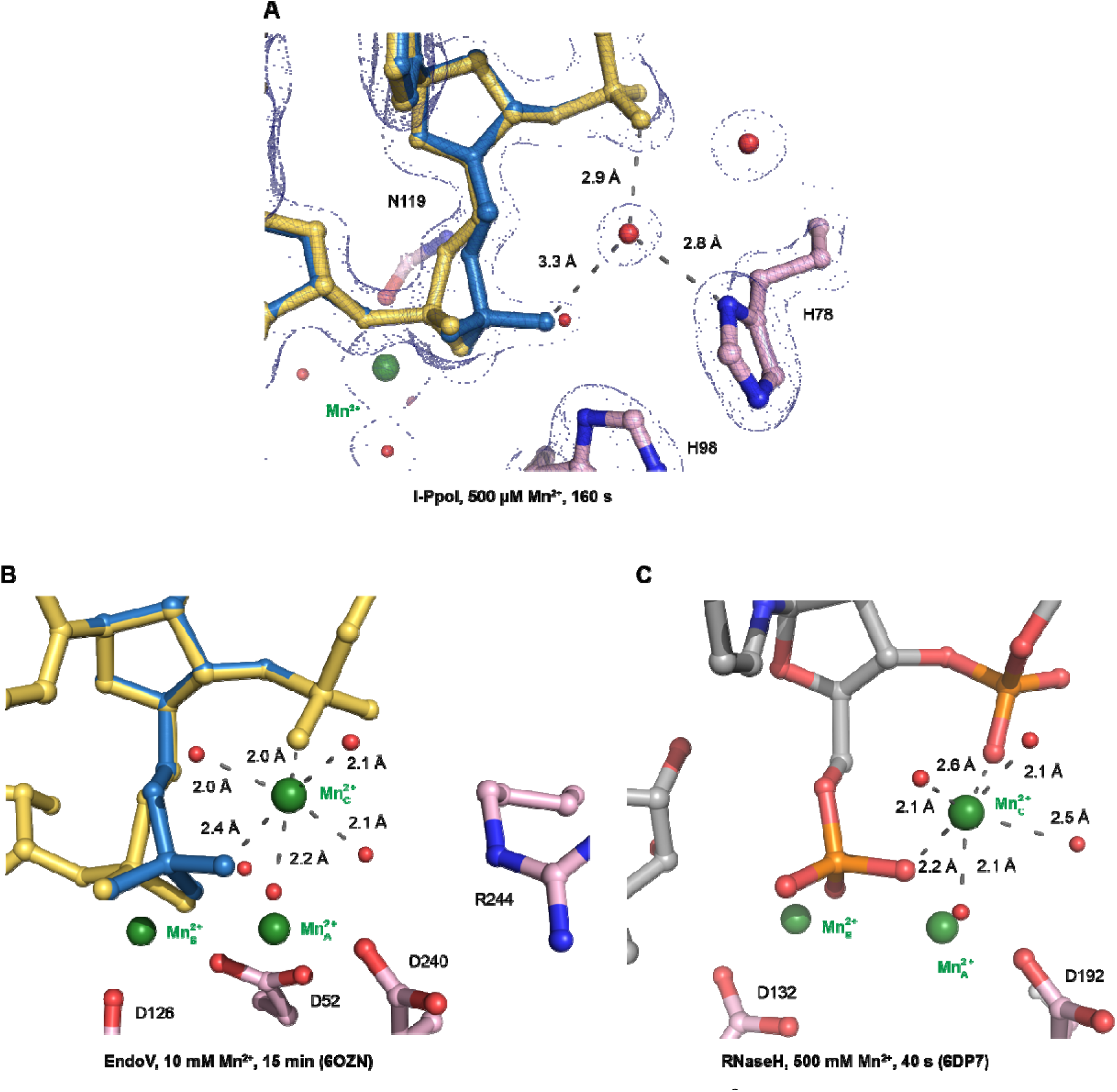
Speculative ligand environment of the transient Me^2+^ in I-PpoI (A) in comparison to EndoV (B) and RNaseH (C). (**A**) The 2F_o_-F_c_ map for Me^2+^, DNA, waters (red spheres), and catalytic residues (blue) was contoured at 2.0 σ. The DNA phosphate conformation within the active site of I-PpoI is looser in comparison to that in EndoV and RNaseH to bind an additional Me^2+^. Carbon atoms of residues within the active site are colored in pink. The Me^2+^ are depicted by green spheres while waters are depicted by red spheres. (**A, B**) the DNA reactant state is colored in yellow while the DNA product state is colored in blue.

**Fig. S6.**
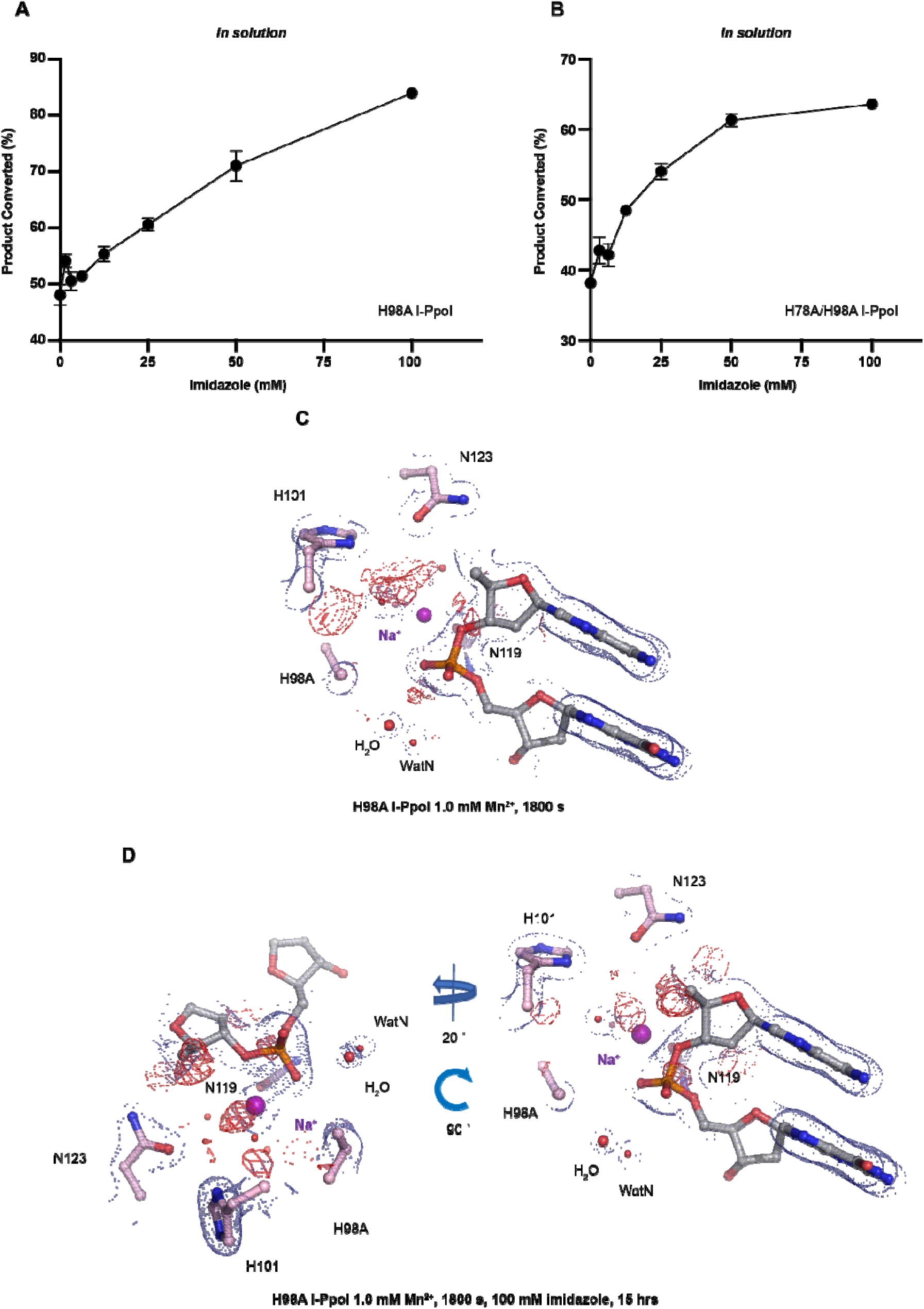
Histidine I-PpoI mutants and partial rescued cleavage activity by imidazole. (**A**) In solution titration of imidazole on DNA cleavage activity by H98A I-PpoI. (**B**) In solution titration of imidazole on DNA cleavage activity by H78A/H98A I-PpoI. (**A, B**) The errors bars represent the standard deviation while the points represent the mean of duplicate measurements for cleaved DNA product. (**C**) Structure of H98A I-PpoI active site after 1 mM Mn^2+^ soaking for 1800 s. (**D**) Structure of H98A I-PpoI active site after 100 mM imidazole for 15 hrs following 1 mM Mn^2+^ soaking for 1800 s. The anomalous map for Mn^2+^ was contoured at 2.0 σ. (**A, D**) The 2F_o_-F_c_ map for Me^2+^, DNA, waters (red spheres), and catalytic residues (blue) was contoured at 2.0 σ.

**Fig. S7.**
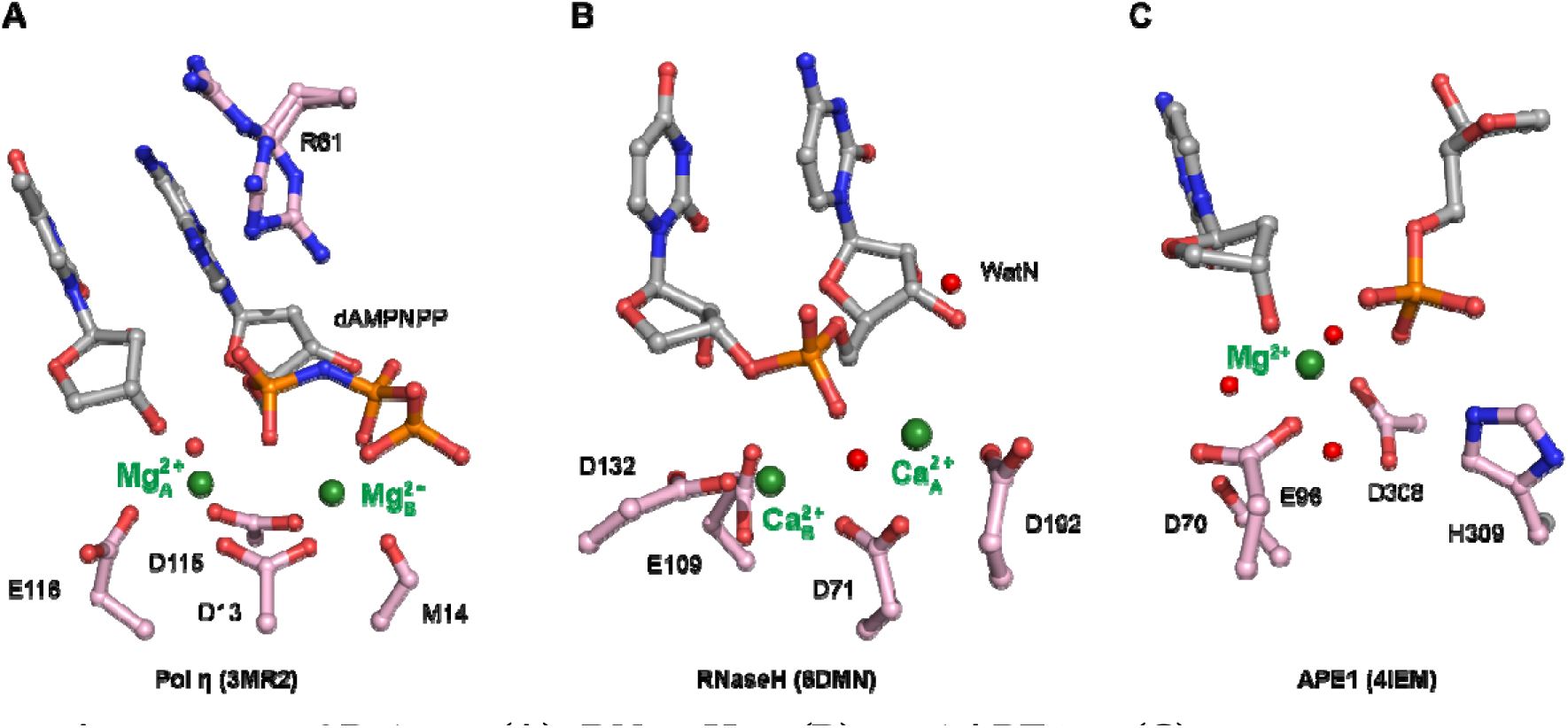
Active sites of Pol η in (A), RNaseH in (B), and APE1 in (C). Carbon atoms of residues within the active site are colored in pink. The Me^2+^ are depicted by green spheres while waters are depicted by red spheres.

**Table S1.**
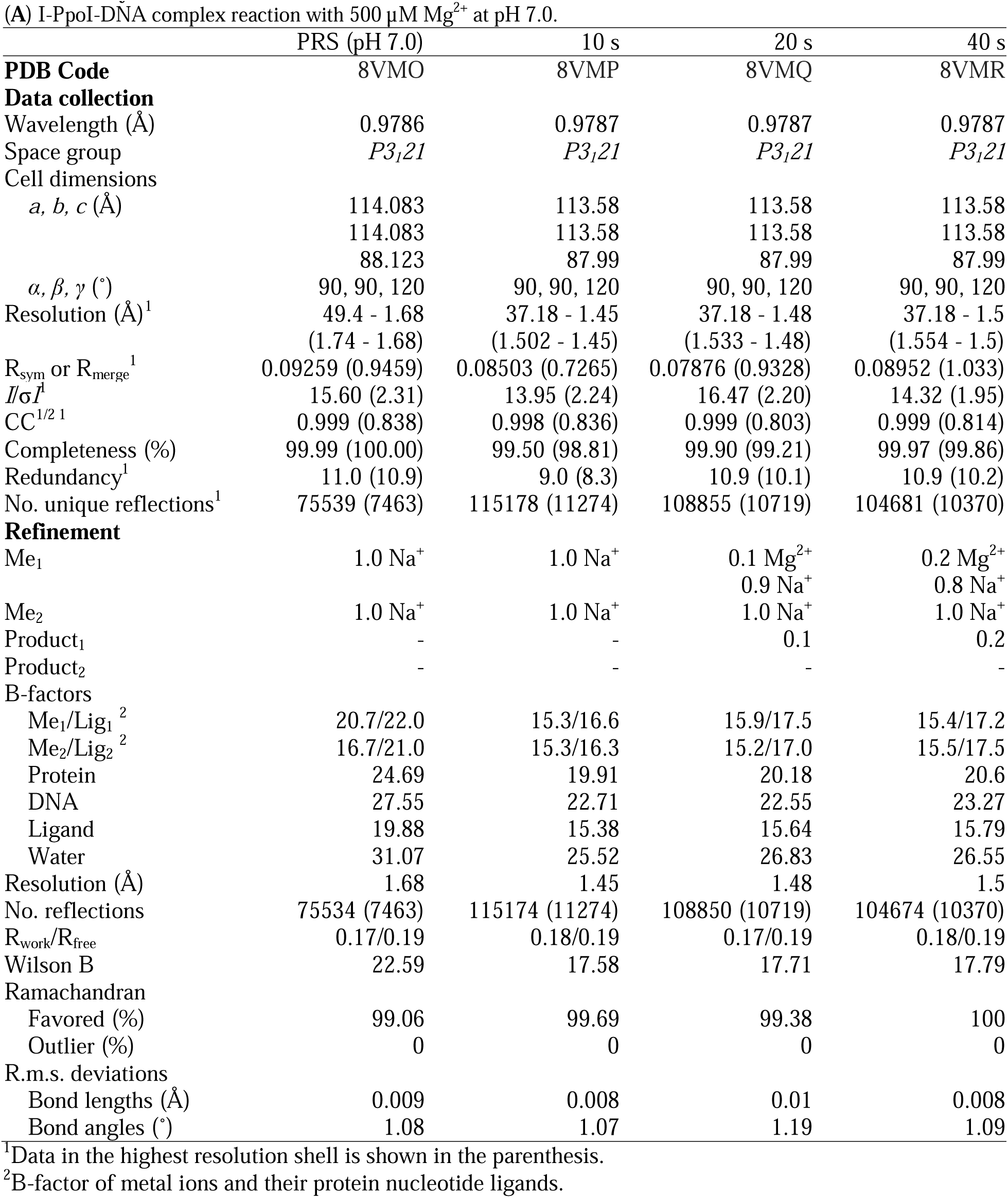
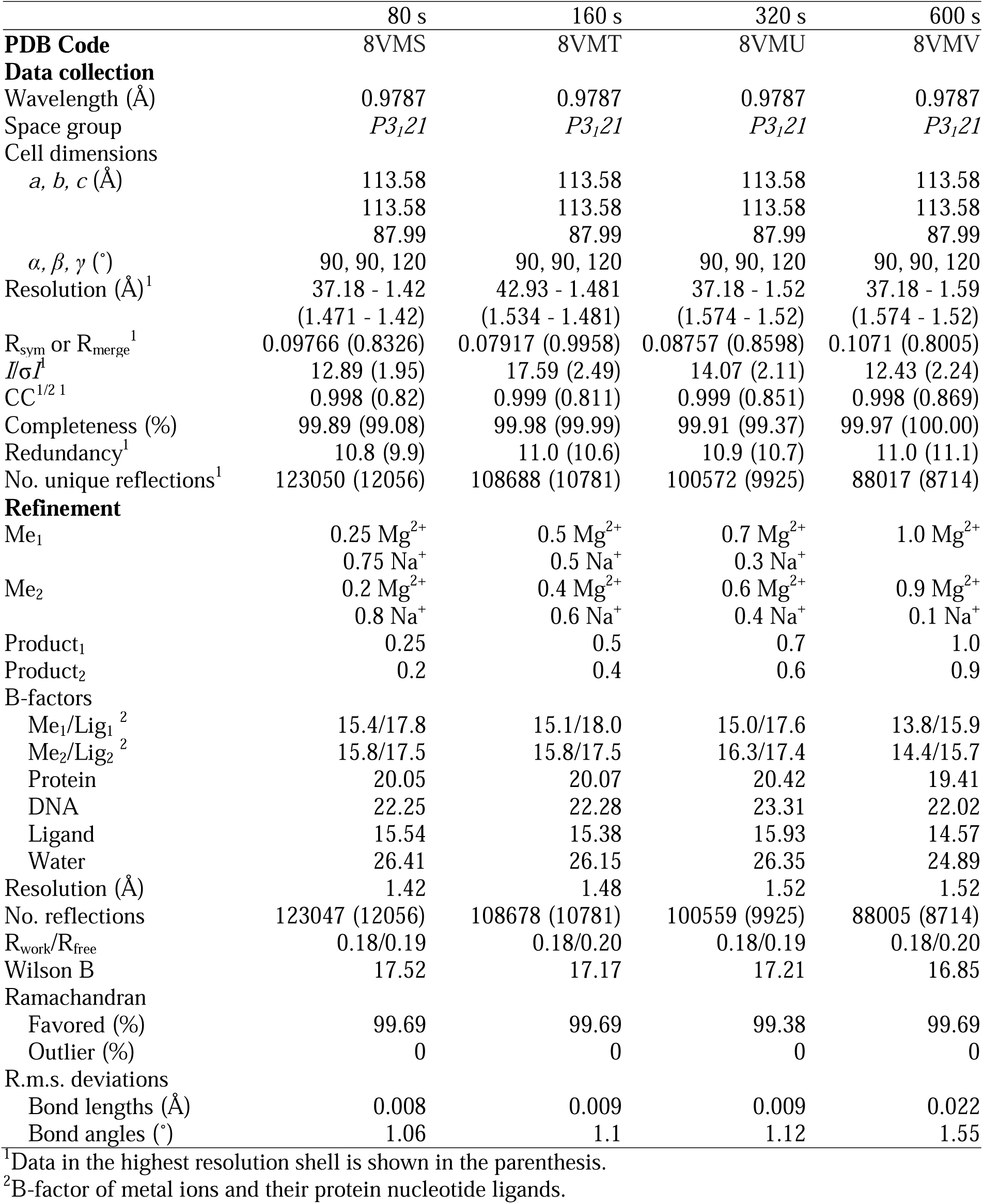
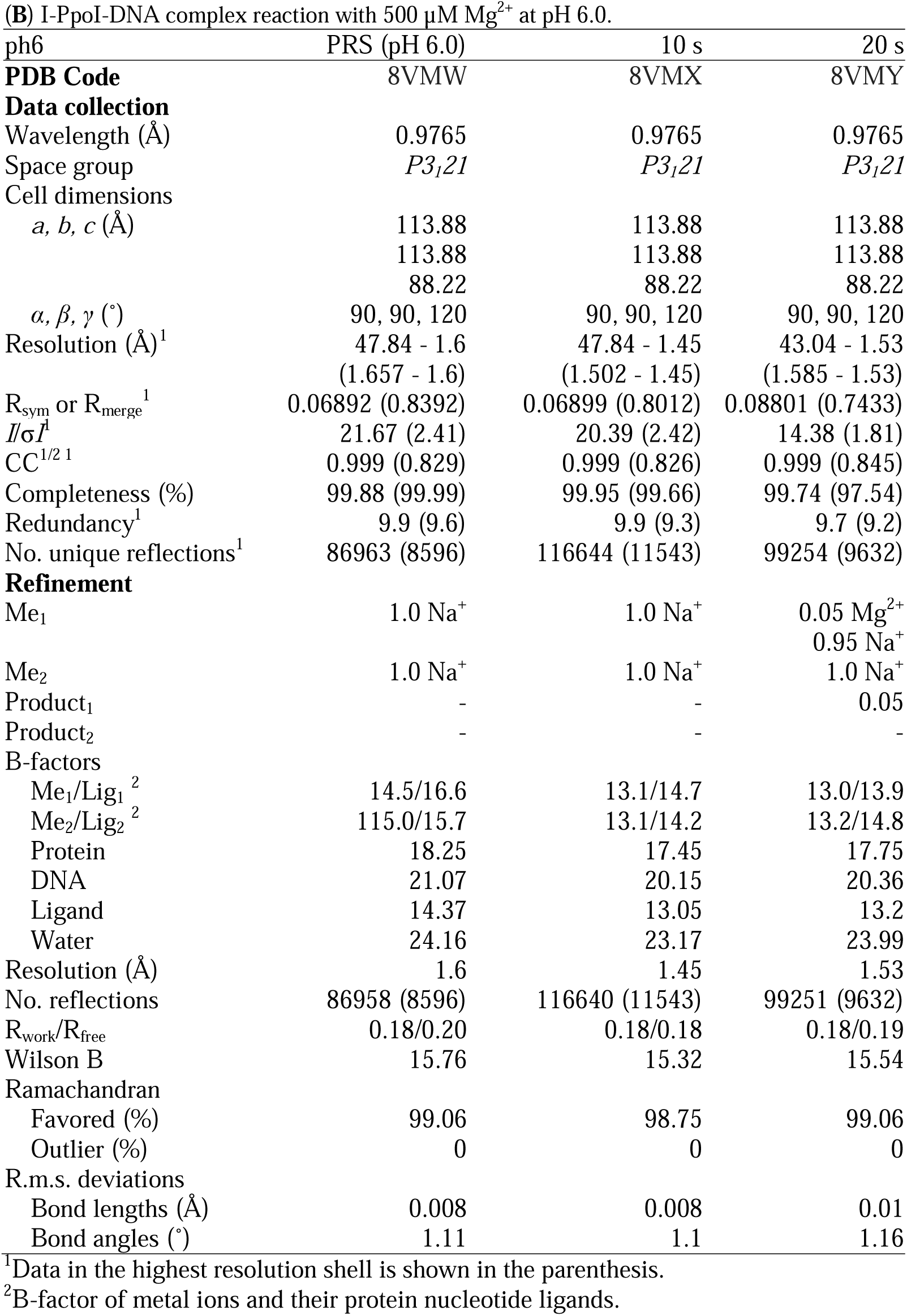
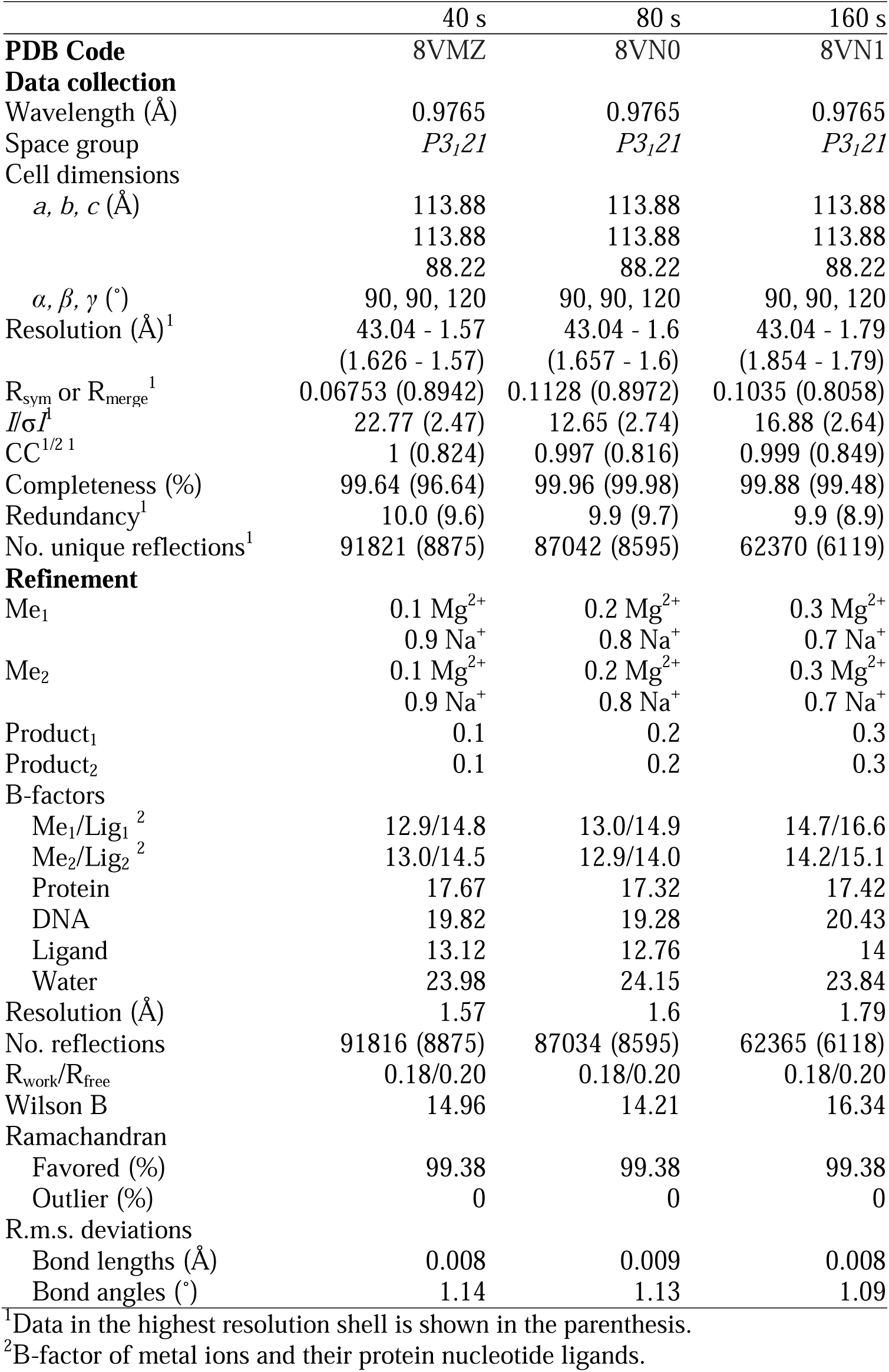
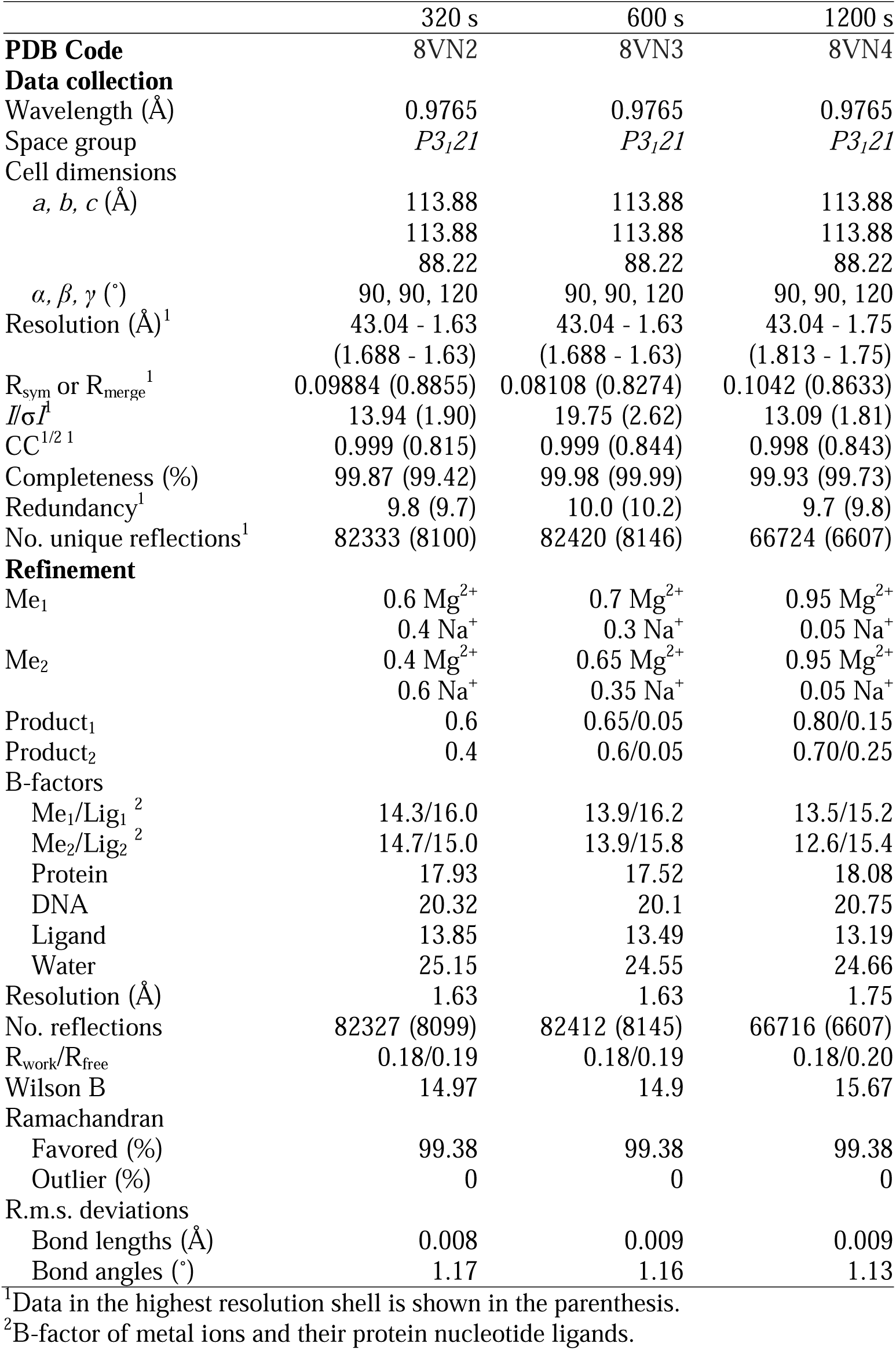
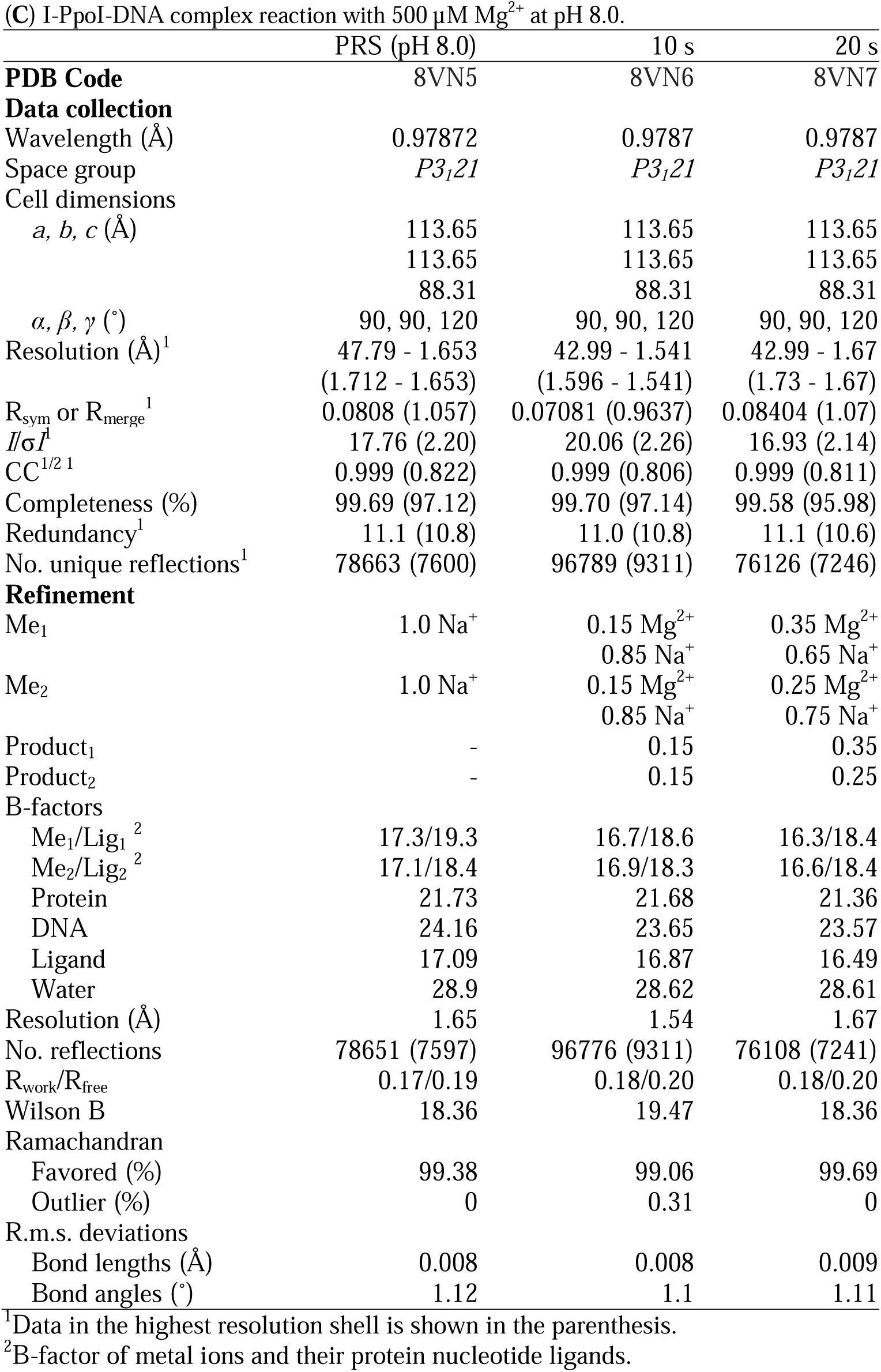
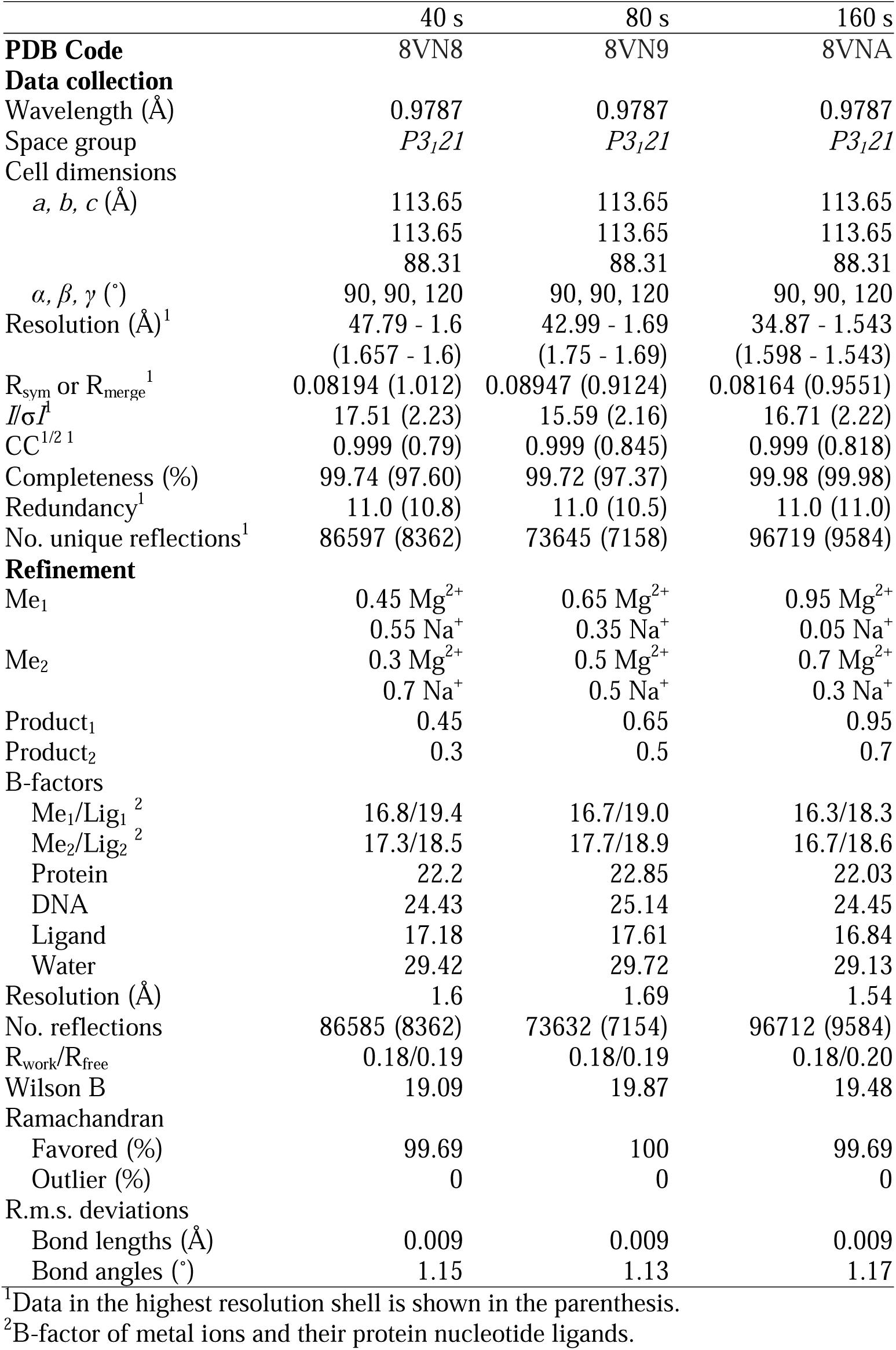
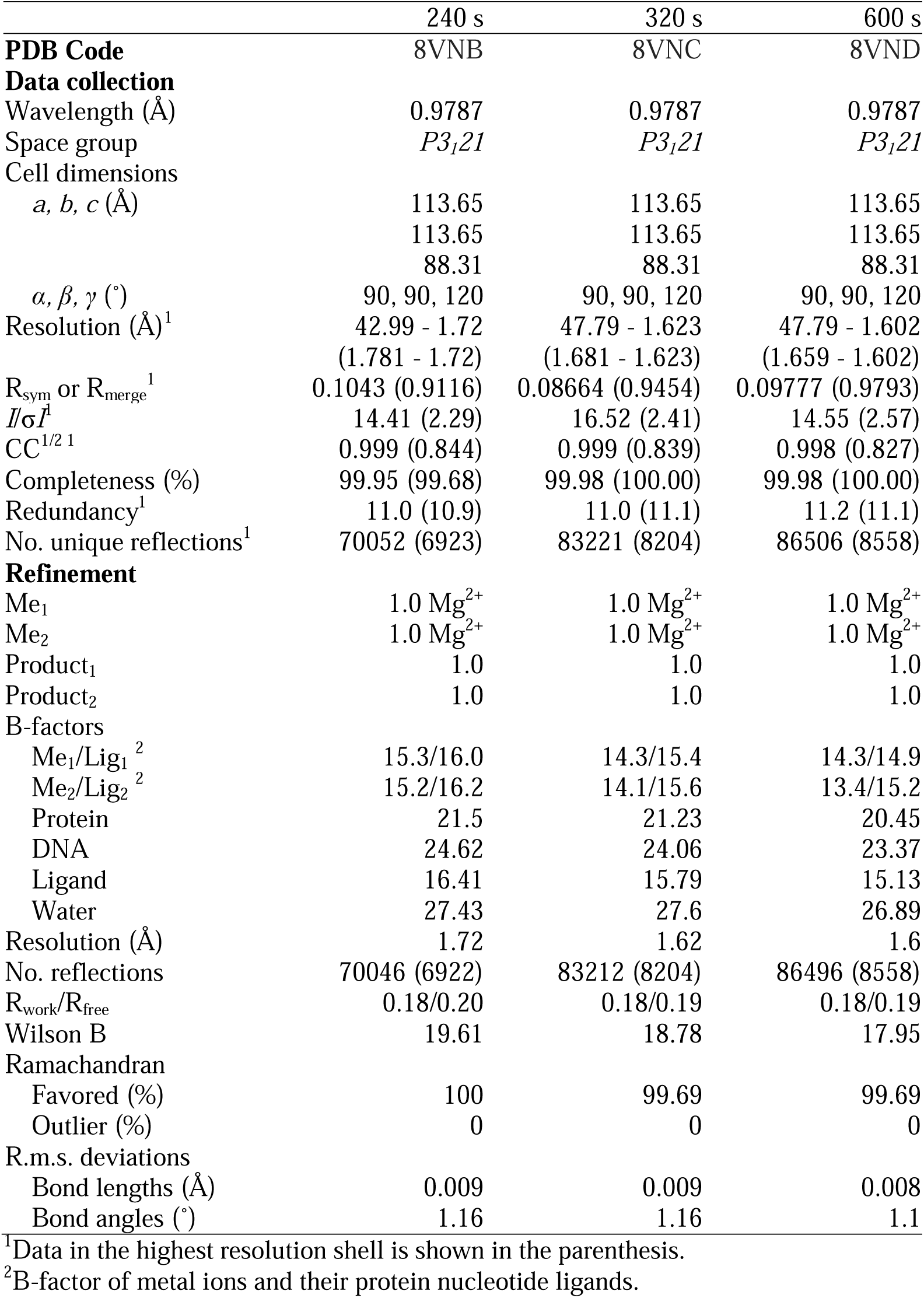
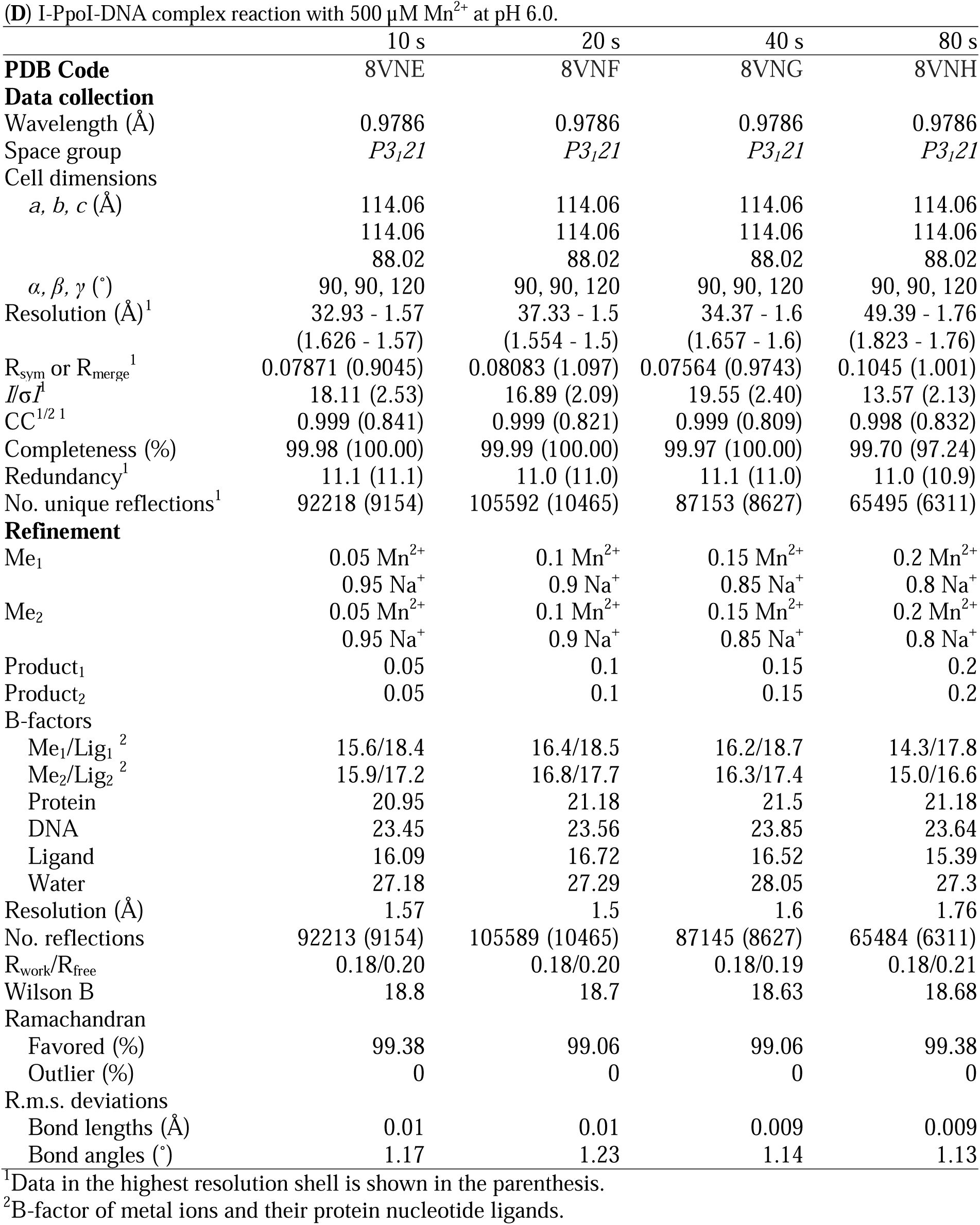
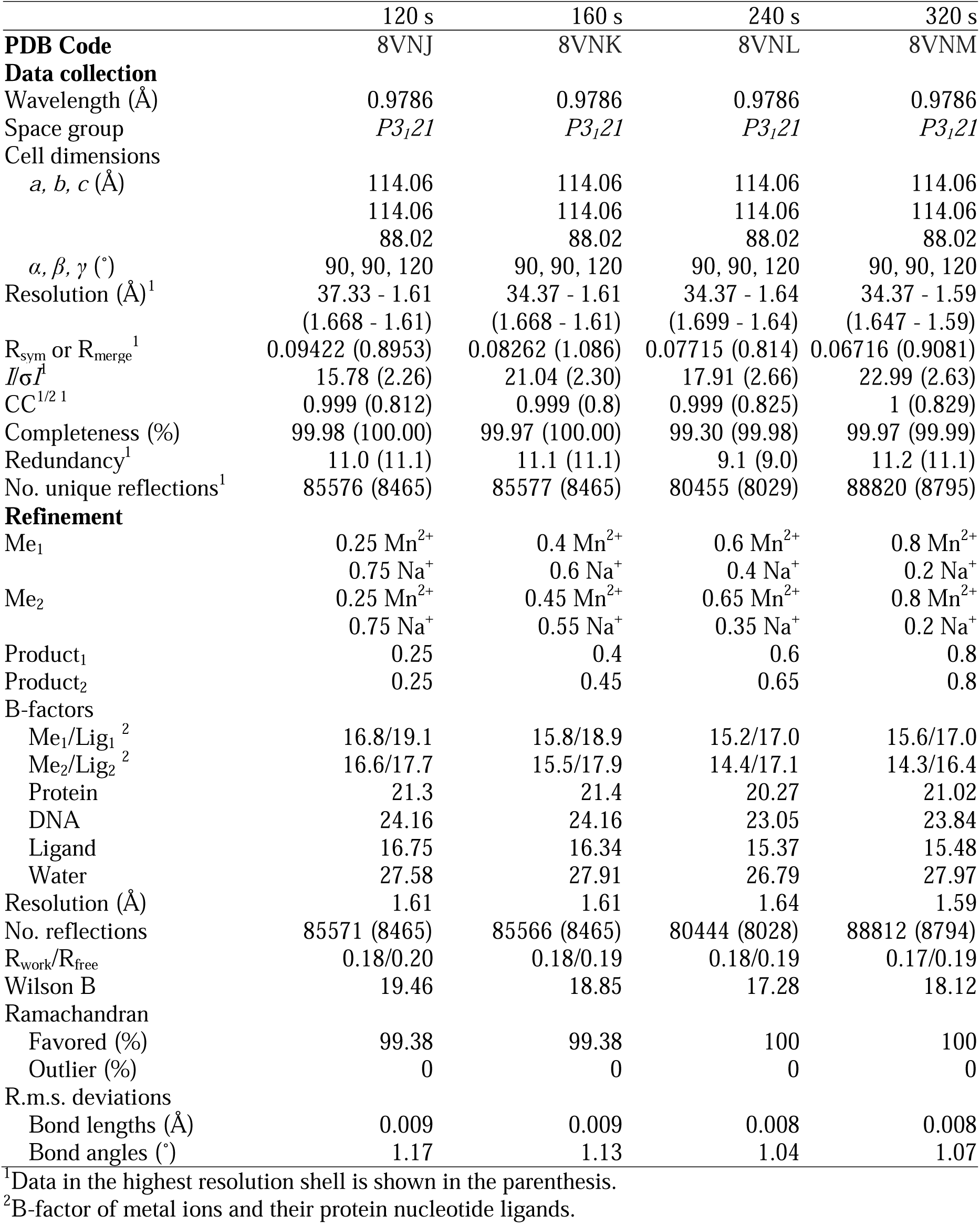
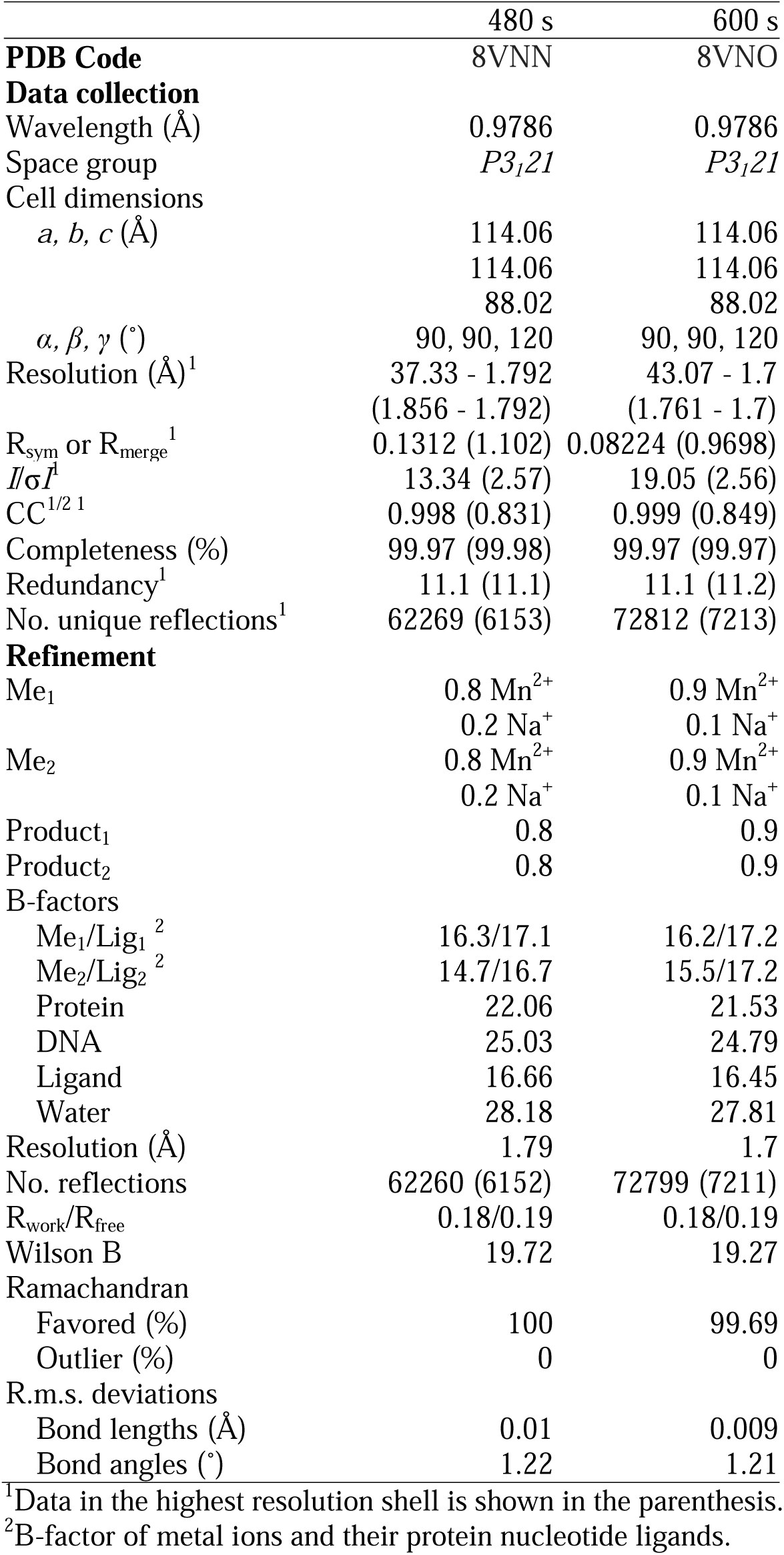
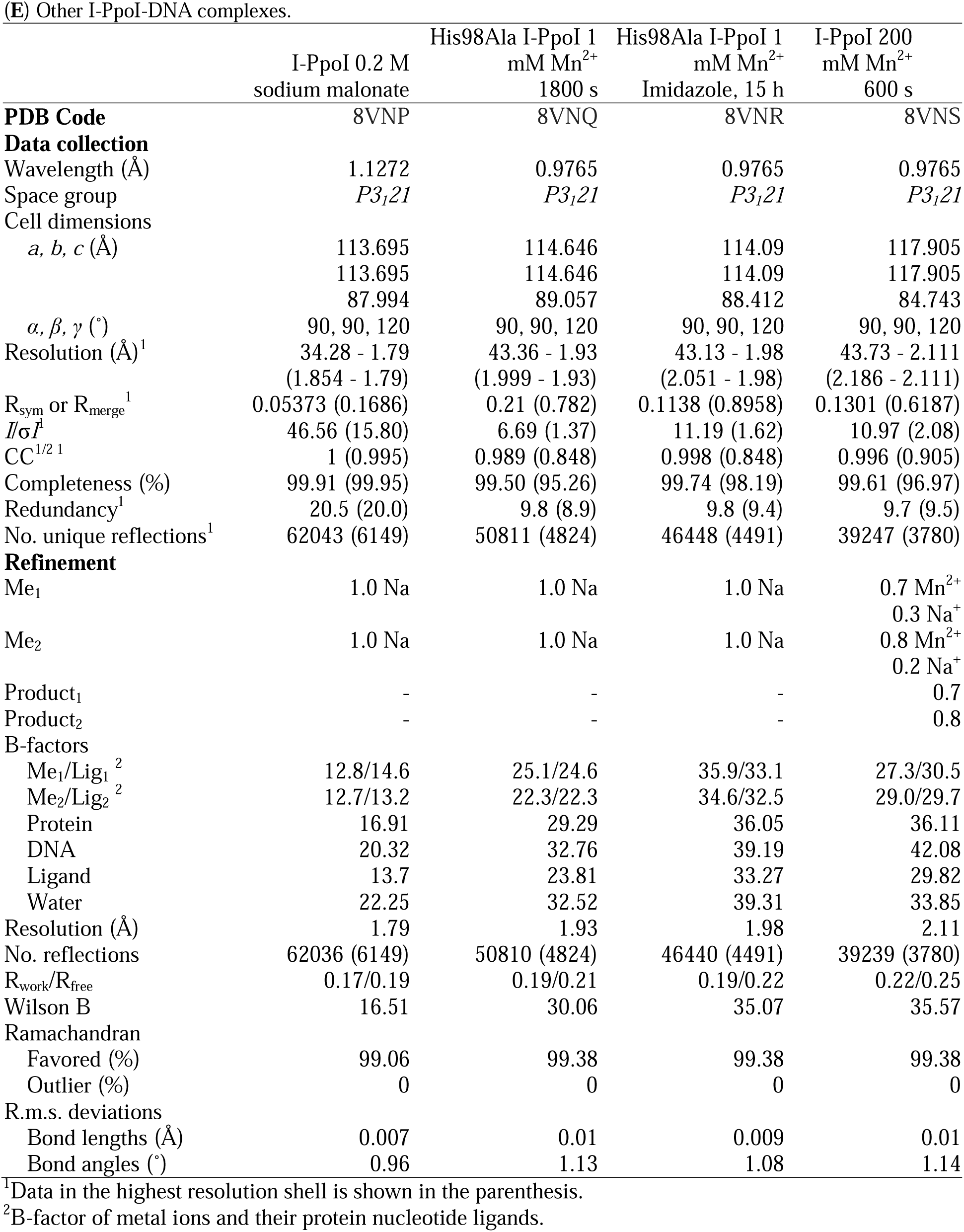
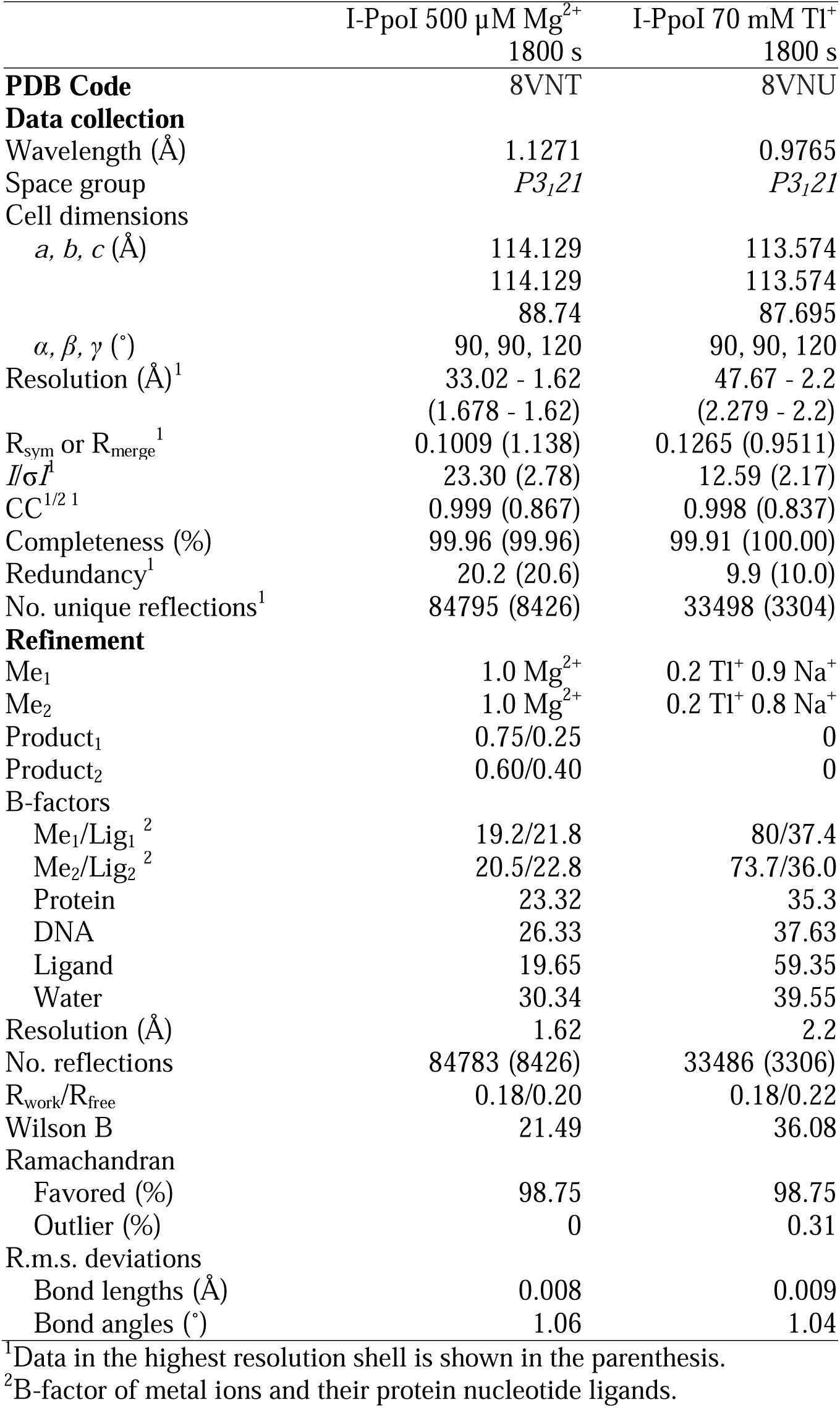
Crystal Diffraction and refinement data.

## References

1 Kao, H.-I. & Bambara, R. A. The Protein Components and Mechanism of Eukaryotic Okazaki Fragment Maturation. Critical Reviews in Biochemistry and Molecular Biology 38, 433–452, doi:10.1080/10409230390259382 (2003).

2 Shen, B. et al. Multiple but dissectible functions of FEN-1 nucleases in nucleic acid processing, genome stability and diseases. Bioessays 27, 717–729, doi:10.1002/bies.20255 (2005).

3 Marti, T. M. & Fleck, O. DNA repair nucleases. Cellular and Molecular Life Sciences CMLS 61, 336–354, doi:10.1007/s00018-003-3223-4 (2004).

4 Mimitou, E. P. & Symington, L. S. DNA end resection: many nucleases make light work. DNA Repair (Amst) 8, 983–995, doi:10.1016/j.dnarep.2009.04.017 (2009).

5 Patel, A. A. & Steitz, J. A. Splicing double: insights from the second spliceosome. Nature Reviews Molecular Cell Biology 4, 960–970, doi:10.1038/nrm1259 (2003).

6 Abelson, J., Trotta, C. R. & Li, H. tRNA splicing. J Biol Chem 273, 12685–12688, doi:10.1074/jbc.273.21.12685 (1998).

7 Chu, C. Y. & Rana, T. M. Small RNAs: regulators and guardians of the genome. J Cell Physiol 213, 412–419, doi:10.1002/jcp.21230 (2007).

8 Moore, M. J. & Proudfoot, N. J. Pre-mRNA processing reaches back to transcription and ahead to translation. Cell 136, 688–700, doi:10.1016/j.cell.2009.02.001 (2009).

9 James, R., Kleanthous, C. & Moore, G. R. The biology of E colicins: paradigms and paradoxes. Microbiology (Reading) 142 **( Pt** **7**), 1569–1580, doi:10.1099/13500872-142-7-1569 (1996).

10 Sorek, R., Kunin, V. & Hugenholtz, P. CRISPR — a widespread system that provides acquired resistance against phages in bacteria and archaea. Nature Reviews Microbiology 6, 181–186, doi:10.1038/nrmicro1793 (2008).

11 Tock, M. R. & Dryden, D. T. The biology of restriction and anti-restriction. Curr Opin Microbiol 8, 466–472, doi:10.1016/j.mib.2005.06.003 (2005).

12 Adli, M. The CRISPR tool kit for genome editing and beyond. Nat Commun 9, 1911, doi:10.1038/s41467-018-04252-2 (2018).

13 Carroll, D. Genome Engineering with Targetable Nucleases. Annu. Rev. Biochem. 83, 409–439, doi:10.1146/annurev-biochem-060713-035418 (2014).

14 Grindley, N. D. F., Whiteson, K. L. & Rice, P. A. Mechanisms of Site-Specific Recombination. Annu. Rev. Biochem. 75, 567–605, doi:10.1146/annurev.biochem.73.011303.073908 (2006).

15 Yang, W. Nucleases: diversity of structure, function and mechanism. Q Rev Biophys 44, 1–93, doi:10.1017/S0033583510000181 (2011).

16 Chen, Y., Gao, T., Wang, Y. & Yang, G. Investigating the Influence of Magnesium Ions on p53-DNA Binding Using Atomic Force Microscopy. Int J Mol Sci 18, doi:10.3390/ijms18071585 (2017).

17 Yang, W. An equivalent metal ion in one- and two-metal-ion catalysis. Nat Struct Mol Biol 15, 1228–1231, doi:10.1038/nsmb.1502 (2008).

18 Samara, N. L. & Yang, W. Cation trafficking propels RNA hydrolysis. Nat Struct Mol Biol 25, 715–721, doi:10.1038/s41594-018-0099-4 (2018).

19 Wu, J., Samara, N. L., Kuraoka, I. & Yang, W. Evolution of Inosine-Specific Endonuclease V from Bacterial DNase to Eukaryotic RNase. Mol Cell 76, 44–56.e43, doi:10.1016/j.molcel.2019.06.046 (2019).

20 Freudenthal, B. D., Beard, W. A., Cuneo, M. J., Dyrkheeva, N. S. & Wilson, S. H. Capturing snapshots of APE1 processing DNA damage. Nat Struct Mol Biol 22, 924–931, doi:10.1038/nsmb.3105 (2015).

21 Whitaker, A. M., Flynn, T. S. & Freudenthal, B. D. Molecular snapshots of APE1 proofreading mismatches and removing DNA damage. Nat Commun 9, 399, doi:10.1038/s41467-017-02175-y (2018).

22 Dupureur, C. M. One is enough: insights into the two-metal ion nuclease mechanism from global analysis and computational studies. Metallomics 2, 609–620, doi:10.1039/C0MT00013B (2010).

23 Garcin, E. D. et al. DNA apurinic-apyrimidinic site binding and excision by endonuclease IV. Nat Struct Mol Biol 15, 515–522, doi:10.1038/nsmb.1414 (2008).

24 Arnoult, D. et al. Mitochondrial release of AIF and EndoG requires caspase activation downstream of Bax/Bak mediated permeabilization. The EMBO Journal 22, 4385–4399, doi: 10.1093/emboj/cdg423 (2003).

25 Lin, J. L. J. et al. Oxidative Stress Impairs Cell Death by Repressing the Nuclease Activity of Mitochondrial Endonuclease G. Cell Reports 16, 279–287, doi: 10.1016/j.celrep.2016.05.090 (2016).

26 Cheng, Y. S., Hsia, K. C., Doudeva, L. G., Chak, K. F. & Yuan, H. S. The crystal structure of the nuclease domain of colicin E7 suggests a mechanism for binding to double-stranded DNA by the H-N-H endonucleases. J Mol Biol 324, 227–236, doi:10.1016/s0022-2836(02)01092-6 (2002).

27 Hsia, K. C. et al. DNA binding and degradation by the HNH protein ColE7. Structure 12, 205–214, doi:10.1016/j.str.2004.01.004 (2004).

28 Doudna, J. A. & Charpentier, E. The new frontier of genome engineering with CRISPR- Cas9. Science 346, 1258096, doi:doi:10.1126/science.1258096 (2014).

29 Ran, F. A. et al. Genome engineering using the CRISPR-Cas9 system. Nature Protocols 8, 2281–2308, doi:10.1038/nprot.2013.143 (2013).

30 Chevalier, B. S. & Stoddard, B. L. Homing endonucleases: structural and functional insight into the catalysts of intron/intein mobility. Nucleic Acids Res 29, 3757–3774, doi:10.1093/nar/29.18.3757 (2001).

31 Stoddard, B. L. Homing endonuclease structure and function. Q Rev Biophys 38, 49–95, doi:10.1017/S0033583505004063 (2005).

32 Wu, C. C., Lin, J. L. J. & Yuan, H. S. Structures, Mechanisms, and Functions of His-Me Finger Nucleases. Trends Biochem Sci 45, 935–946, doi:10.1016/j.tibs.2020.07.002 (2020).

33 Galburt, E. A. et al. A novel endonuclease mechanism directly visualized for I-PpoI. Nat Struct Biol 6, 1096–1099, doi:10.1038/70027 (1999).

34 Pommer, A. J. et al. Mechanism and cleavage specificity of the H-N-H endonuclease colicin E911Edited by J. Karn. Journal of Molecular Biology 314, 735–749, doi:10.1006/jmbi.2001.5189 (2001).

35 Li, C. L. et al. DNA binding and cleavage by the periplasmic nuclease Vvn: a novel structure with a known active site. The EMBO Journal 22, 4014–4025, doi:10.1093/emboj/cdg377 (2003).

36 Shen, B. W., Landthaler, M., Shub, D. A. & Stoddard, B. L. DNA Binding and Cleavage by the HNH Homing Endonuclease I-HmuI. Journal of Molecular Biology 342, 43–56, doi:10.1016/j.jmb.2004.07.032 (2004).

37 Schwank, G. et al. Functional Repair of CFTR by CRISPR/Cas9 in Intestinal Stem Cell Organoids of Cystic Fibrosis Patients. Cell Stem Cell 13, 653–658, doi:10.1016/j.stem.2013.11.002 (2013).

38 Sharma, G., Sharma, A. R., Bhattacharya, M., Lee, S.-S. & Chakraborty, C. CRISPR- Cas9: A Preclinical and Clinical Perspective for the Treatment of Human Diseases. Molecular Therapy 29, 571–586, doi:10.1016/j.ymthe.2020.09.028 (2021).

39 Khan, F. A. et al. CRISPR/Cas9 therapeutics: a cure for cancer and other genetic diseases. Oncotarget 7, 52541–52552, doi:10.18632/oncotarget.9646 (2016).

40 Sun, W. et al. Structures of Neisseria meningitidis Cas9 Complexes in Catalytically Poised and Anti-CRISPR-Inhibited States. Mol Cell 76, 938–952.e935, doi:10.1016/j.molcel.2019.09.025 (2019).

41 Zhang, Y. et al. Catalytic-state structure and engineering of Streptococcus thermophilus Cas9. Nature Catalysis 3, 813–823, doi:10.1038/s41929-020-00506-9 (2020).

42 Zhu, X. et al. Cryo-EM structures reveal coordinated domain motions that govern DNA cleavage by Cas9. Nat Struct Mol Biol 26, 679–685, doi:10.1038/s41594-019-0258-2 (2019).

43 Das, A. et al. Coupled catalytic states and the role of metal coordination in Cas9. Nature Catalysis 6, 969–977, doi:10.1038/s41929-023-01031-1 (2023).

44 Flick, K. E., Jurica, M. S., Monnat, R. J. & Stoddard, B. L. DNA binding and cleavage by the nuclear intron-encoded homing endonuclease I-PpoI. Nature 394, 96–101, doi:10.1038/27952 (1998).

45 Freudenthal, Bret D., Beard, William A., Shock, David D. & Wilson, Samuel H. Observing a DNA Polymerase Choose Right from Wrong. Cell 154, 157–168, doi:10.1016/j.cell.2013.05.048 10.1016/j.cell.2013.05.048 (2013).

46 Gao, Y. & Yang, W. Capture of a third Mg2+ is essential for catalyzing DNA synthesis. Science 352, 1334–1337, doi:10.1126/science.aad9633 (2016).

47 Nakamura, T., Zhao, Y., Yamagata, Y., Hua, Y.-j. & Yang, W. Watching DNA polymerase η make a phosphodiester bond. Nature 487, 196–201, doi:10.1038/nature11181 (2012).

48 Vyas, R., Reed, A. J., Tokarsky, E. J. & Suo, Z. Viewing Human DNA Polymerase β Faithfully and Unfaithfully Bypass an Oxidative Lesion by Time-Dependent Crystallography. J. Am. Chem. Soc. 137, 5225–5230, doi:10.1021/jacs.5b02109 (2015).

49 Chim, N., Meza, R. A., Trinh, A. M., Yang, K. & Chaput, J. C. Following replicative DNA synthesis by time-resolved X-ray crystallography. Nat Commun 12, 2641, doi:10.1038/s41467-021-22937-z (2021).

50 Gregory, M. T., Gao, Y., Cui, Q. & Yang, W. Multiple deprotonation paths of the nucleophile 3′-OH in the DNA synthesis reaction. Proceedings of the National Academy of Sciences 118, e2103990118, doi:10.1073/pnas.2103990118 (2021).

51 Demir, M. et al. Structural snapshots of base excision by the cancer-associated variant MutY N146S reveal a retaining mechanism. Nucleic Acids Research 51, 1034–1049, doi:10.1093/nar/gkac1246 (2023).

52 Deerfield, D. W., Fox, D. J., Head-Gordon, M., Hiskey, R. G. & Pedersen, L. G. Interaction of calcium and magnesium ions with malonate and the role of the waters of hydration: a quantum mechanical study. J. Am. Chem. Soc. 113, 1892–1899, doi:10.1021/ja00006a004 (1991).

53 Cowan, J. A. Structural and catalytic chemistry of magnesium-dependent enzymes. Biometals 15, 225–235, doi:10.1023/A:1016022730880 (2002).

54 Auffinger, P., D’Ascenzo, L. & Ennifar, E. in The Alkali Metal Ions: Their Role for Life (eds Astrid Sigel, Helmut Sigel, & Roland K. O. Sigel) 167–201 (Springer International Publishing, 2016).

55 Kiser, P. D., Lorimer, G. H. & Palczewski, K. Use of thallium to identify monovalent cation binding sites in GroEL. Acta Crystallogr Sect F Struct Biol Cryst Commun 65, 967–971, doi:10.1107/s1744309109032928 (2009).

56 Eastberg, J. H., Eklund, J., Monnat, R. & Stoddard, B. L. Mutability of an HNH Nuclease Imidazole General Base and Exchange of a Deprotonation Mechanism. Biochemistry 46, 7215–7225, doi:10.1021/bi700418d (2007).

57 Furuhata, Y. & Kato, Y. Asymmetric Roles of Two Histidine Residues in Streptococcus pyogenes Cas9 Catalytic Domains upon Chemical Rescue. Biochemistry 60, 194–200, doi:10.1021/acs.biochem.0c00766 (2021).

58 Moon, A. F. et al. Structural insights into catalytic and substrate binding mechanisms of the strategic EndA nuclease from Streptococcus pneumoniae. Nucleic Acids Research 39, 2943–2953, doi:10.1093/nar/gkq1152 (2010).

59 Maghsoud, Y., Jayasinghe-Arachchige, V. M., Kumari, P., Cisneros, G. A. & Liu, J. Leveraging QM/MM and Molecular Dynamics Simulations to Decipher the Reaction Mechanism of the Cas9 HNH Domain to Investigate Off-Target Effects. Journal of Chemical Information and Modeling 63, 6834–6850, doi:10.1021/acs.jcim.3c01284 (2023).

60 Wilson, M. A. Mapping Enzyme Landscapes by Time-Resolved Crystallography with Synchrotron and X-Ray Free Electron Laser Light. Annu Rev Biophys 51, 79–98, doi:10.1146/annurev-biophys-100421-110959 (2022).

61 Branden, G. & Neutze, R. Advances and challenges in time-resolved macromolecular crystallography. Science 373, doi:10.1126/science.aba0954 (2021).

62 Maestre-Reyna, M. et al. Visualizing the DNA repair process by a photolyase at atomic resolution. Science 382, eadd7795, doi:doi:10.1126/science.add7795 (2023).

63 Christou, N.-E. et al. Time-resolved crystallography captures light-driven DNA repair. Science 382, 1015–1020, doi:doi:10.1126/science.adj4270 (2023).

64 Reed, A. J. & Suo, Z. Time-Dependent Extension from an 8-Oxoguanine Lesion by Human DNA Polymerase Beta. J. Am. Chem. Soc. 139, 9684–9690, doi:10.1021/jacs.7b05048 (2017).

65 Chang, C., Lee Luo, C. & Gao, Y. In crystallo observation of three metal ion promoted DNA polymerase misincorporation. Nat Commun 13, 2346, doi:10.1038/s41467-022-30005-3 (2022).

66 Nakamura, T. & Yamagata, Y. Visualization of mutagenic nucleotide processing by *Escherichia coli* MutT, a Nudix hydrolase. Proceedings of the National Academy of Sciences 119, e2203118119, doi:doi:10.1073/pnas.2203118119 (2022).

67 Tsutakawa, S. E. et al. Conserved Structural Chemistry for Incision Activity in Structurally Non-homologous Apurinic/Apyrimidinic Endonuclease APE1 and Endonuclease IV DNA Repair Enzymes*. Journal of Biological Chemistry 288, 8445–8455, doi:10.1074/jbc.M112.422774 (2013).

68 Galburt, E. A. et al. Conformational Changes and Cleavage by the Homing Endonuclease I-PpoI: A Critical Role for a Leucine Residue in the Active Site. Journal of Molecular Biology 300, 877–887, doi:10.1006/jmbi.2000.3874 (2000).

69 Nierzwicki, Ł. et al. Principles of target DNA cleavage and the role of Mg2+ in the catalysis of CRISPR-Cas9. Nat Catal 5, 912–922, doi:10.1038/s41929-022-00848-6 (2022).

70 Ashworth, J. et al. Computational redesign of endonuclease DNA binding and cleavage specificity. Nature 441, 656–659, doi:10.1038/nature04818 (2006).

71 Kabsch, W. XDS. Acta crystallographica. Section D, Biological crystallography 66, 125–132, doi:10.1107/S0907444909047337 (2010).

72 Adams, P. D. et al. PHENIX: a comprehensive Python-based system for macromolecular structure solution. Acta Crystallogr D Biol Crystallogr 66, 213–221, doi:10.1107/s0907444909052925 (2010).

73 Emsley, P., Lohkamp, B., Scott, W. G. & Cowtan, K. Features and development of Coot. Acta Crystallogr D Biol Crystallogr 66, 486–501, doi:10.1107/s0907444910007493 (2010).

74 Morin, A. et al. Collaboration gets the most out of software. eLife 2, e01456, doi:10.7554/eLife.01456 (2013).

